# Historical forest disturbance results in variation in functional resilience of seed dispersal mutualisms

**DOI:** 10.1101/2022.05.28.493853

**Authors:** Carmela M. Buono, Jesse Lofaso, Will Smisko, Carly Gerth, John Santare, Kirsten M. Prior

**Affiliations:** Binghamton University, Department of Biological Sciences, NY, United States

**Author notes:** **Corresponding author:** Carmela M. Buono.

**Keywords:** land use change, seed dispersal, mutualism, myrmecochory, resilience, ecosystem function, *Aphaenogaster*, *Arion*, ants, animal–plant interaction, deciduous forests

## Abstract

Mutualistic interactions provide essential ecosystem functions, such as promoting and maintaining diversity. Understanding if functionally important mutualisms are resilient (able to resist and recover) to anthropogenic disturbance is important to understand the capacity for diversity to recover. Animal-mediated seed dispersal supports plant population growth and community structure, and disturbance of this function can threaten plant diversity and contribute to low resiliency. Ant-mediated seed dispersal mutualisms are particularly sensitive to anthropogenic disturbance, as they rely on one to a few high-quality dispersal partners. In North American eastern deciduous forests (NAEDF), ants in the genus *Aphaenogaster* are “keystone dispersers” of 30-40% of understory forbs adapted to dispersal by ants (myrmecochores). The majority of present day NAEDF have regenerated from previous disturbance in the form of historical land use change (HLUC), due to clearing for agriculture. Previous studies have revealed that myrmecochore diversity is not resilient to HLUC. Here, we ask if seed dispersal mutualisms are resilient to HLUC and if decreases in mutualistic interactions with partners, *Aphaenogaster* sp., or increases in antagonistic interactions cause degradation of function. In a large-scale natural experiment (20 sites), we measured seed removal, the abundance of mutualistic partners and other invertebrates interacting with seeds, myrmecochore cover and diversity, along with ant habitat and forest structure. We found lower and more variable seed removal in secondary forests compared to remnant forests. A path analysis of all forests revealed that abundance of mutualists was the primary determinant of variation in seed removal, and that seed damage by antagonists (invasive slugs) negatively affected dispersal and was higher in secondary forests. In a path analysis of remnant forests, the link between mutualist abundance and seed removal was absent, but present in the secondary forest path, suggesting that seed dispersal is more variable and dependent on mutualist abundance in secondary forests and is stable and high in remnant forests. Here we show that functional resilience to HLUC is variable and may impede recovery of understory plant communities. This work provides key insights on the effects of anthropogenic disturbance on mutualistic interactions and how the resilience of critical ecosystem functions impacts diversity resiliency.

## Introduction

Mutualistic interactions provide essential ecosystem functions and contribute to the maintenance of diversity in ecosystems (Goheen & Palmer 2010; Prior et al. 2015; Kaiser-Bunbury et al. 2017; Wandrag et al. 2017). Understanding if mutualistic interactions are functionally resilient (able to resist and recover) to anthropogenic disturbances will reveal if diversity benefitting from interactions has the capacity for recovery (Oliver et al. 2015; Kaiser-Bunbury et al. 2017; Rogers et al. 2017). Shifts in anthropogenic land use leave ecosystems to naturally regenerate from wide-scale disturbance, and variation in legacies may prevent or alter recovery trajectories, leading to reduced or altered interactions and diversity (Holling 1973; Suding et al. 2004; Sabatini et al. 2014; Kaiser-Bunbury et al. 2017). Whether or not functionally important mutualistic interactions are resilient to anthropogenic disturbance is an open question necessary to uncover the capacity for recovery of diversity in ecosystems.

Animal-mediated seed dispersal is a functionally important mutualism (Schupp 1993; Ness et al. 2009; Rogers et al. 2017; Wandrag et al. 2017), where plants benefit from having a dispersal agent that increases dispersal distance from maternal plants, protection from seed predators, and directed dispersal to more favorable location (Tiffney & Mazer 1995; Kalisz et al. 1999; Wenny 2001; Bronstein et al. 2006; Giladi 2006). Animal-mediated seed dispersal benefits restoration, given its positive role in plant population growth and distribution (Wunderle 1997; da Silva et al. 2015; De Almeida et al. 2020), and uncovering if seed dispersal is resilient post disturbance can better inform restoration planning.

Myrmecochory, or ant-mediated seed dispersal, is a widespread dispersal syndrome, including ∼11,000 plant species worldwide (Lengyel et al. 2009). Myrmecochorous plants are adapted to dispersal by ants, having a lipid-rich appendage (elaiosome) that attracts ants and provides a food reward. Myrmecochory is a particularly specialized diffuse mutualism, with one or a handful of ant species dispersing most seeds (Bronstein et al. 2006; Giladi 2006; Gove et al. 2007; Manzaneda & Rey 2009; Ness et al. 2009; Warren et al. 2014). Asymmetrical diffuse mutualisms are sensitive to anthropogenic disturbance, as changes in the presence, abundance, and interaction of effective animal disperser partners can disrupt interactions (Schupp 1993; Traveset & Richardson 2006; Schleuning et al. 2011; Prior et al. 2015), with cascading impacts on plant communities (Christian 2001; Prior et al. 2015; Rogers et al. 2017).

North American eastern deciduous forests (NAEDF) are a myrmecochory hotspot (Lengyel et al. 2009) with 30-40% of understory forbs possessing elaiosomes (Beattie & Culver 1981; Handel et al. 1981). High-quality dispersers find seeds quickly, do not harm seeds, and deposit seeds outside of nests (Giladi 2006; Canner et al. 2012; Prior et al. 2014; Prior et al. 2015; Gordon et al. 2019; Meadley Dunphy et al. 2020). Poor-quality dispersers interact with seeds but fail to disperse seeds and might damage seeds (Christian 2001; Bronstein et al. 2006; Giladi 2006; Stuble et al. 2011; Warren & Giladi 2014; Parker et al. 2021). In NAEDF, ants belonging to the genus *Aphaenogaster* are considered “keystone” dispersers, responsible for majority of dispersal events (Ness et al. 2009).

NAEDF are impacted by various types of anthropogenic disturbances (Reich & Frelich 2002). Most contemporary forests are secondary, having passively regenerated from historical clearing for agriculture or timber harvesting (Flinn & Vellend 2005; Flinn & Marks 2007). Herbaceous understory communities are not resilient to historical clearing with secondary forests lacking diversity, especially of myrmecochores, (Bellemare et al. 2002; Mitchell et al. 2002; Flinn & Vellend 2005; Griffiths & McGee 2018). Several factors likely contribute to low resiliency, including recruitment limitation to secondary fragments from source populations (Flinn & Vellend 2005). Agricultural disturbances leave legacy effects in soils that impact herbaceous plants such as elevated pH and nutrients and lower organic matter (Koerner et al. 1997; Dyer 2010). Less is known about whether functionally important interactions, such as seed dispersal, are resilient to historical land use change (HLUC) and if reduced function contributes to low understory recovery (except see Mitchell et al. 2002; Kiel et al. 2020; Parker et al. 2021).

Reduction in ant partner presence, abundance, or interactions could slow understory recovery in secondary forests (Mitchell et al. 2002; Kiel et al. 2020; Parker et al. 2021). It is well established that *Aphaenogaster* presence and abundance positively influences seed dispersal, distribution and community structure of myrmecochores (Kalisz et al. 1999; Ness et al. 2009; Warren & Giladi 2014; Prior et al. 2015). One way in which myrmecochory may not be resilient to HLUC is if the keystone mutualist, *Aphaenogaster*, are absent or have lower abundances. Prior clearing or soil disturbance could reduce *Aphaenogaster* abundance, and recovery from populations in intact forests could be limited (Schmidt et al. 2013). Ant abundance in secondary forests could also be negatively impacted by changes to forest floor conditions, including altered canopy structure influencing soil microhabitat (Warren et al. 2012), or altered leaf litter and woody debris, which provides ant nesting habitat (Lubertazzi 2012).

Myrmecochory might also not be resilient to HLUC if there is an increase in organisms that interact antagonistically with *Aphaenogaster* or seeds. *Aphaenogaster* is generally subordinate to other ant species and competition with other ants could reduce *Aphaenogaster* abundance (Ness 2004; Warren et al. 2020; Parker et al. 2021). Other ants in NAEDF are lower quality dispersers that fail to disperse seeds or harm seeds (Ness et al. 2004; Giladi 2006; Rodriguez-Cabal et al. 2012; Parker et al. 2021). If secondary forest conditions favor other ant species, they could disrupt seed dispersal directly by harming seeds, failing to disperse seeds, moving them short distances, or indirectly by competing with *Aphaenogaster* and reducing their effectiveness (Ness 2004; Leal et al. 2015; Parker et al. 2021). Finally, other invertebrate organisms interact with myrmecochore seeds and could affect dispersal (Gunther & Lanza 1989; Jules 1998). For example, in NAEDF, the invasive slug, *Arion subfuscus*, occurs in disturbed forest habitats, such as forest edges, and damage myrmecochorous seeds by robbing elaiosomes and preventing dispersal (Meadley Dunphy et al. 2016; Kiel et al. 2020; Parker et al. 2021).

Here, we examine if seed dispersal mutualisms are resilient to wide-scale disturbances resulting from historical land use change (HLUC). In particular, we ask whether seed dispersal is lower in secondary forests compared to remnant forests, and if so, whether decreases in interactions with mutualistic partners or increases in antagonistic interactions contributes to degradation of function. We conducted a natural experiment in 20 paired remnant and secondary forests, where we compared seed dispersal of myrmecochorous seeds, along with myrmecochore cover and diversity. To uncover how abundances of organisms interacting with seeds affects variation in dispersal, we measured the abundance of the keystone disperser, along with other invertebrates (ants, slugs) interacting with seeds. We also measured several forest structure and ant habitat factors (canopy cover, ant nesting sites, and abiotic factors) and performed a path analysis to reveal how forest conditions affect the abundance of mutualistic and antagonistic organisms interacting with seeds and seed dispersal. We predict that seed dispersal will not be resilient to HLUC and be lower in secondary forests due to lower abundance or fewer interactions with mutualists or increased abundance or interactions with antagonists. We also predict a disruption in seed dispersal mutualisms may relate to lower myrmecochore diversity and cover in secondary forests. Better understanding the effects of anthropogenic disturbance on this critical mutualistic interaction, in particular its effect on mutualists and antagonists, provides key insights into the functional resiliency of mutualisms, and its influences on the resilience of diversity. Understanding resiliency of seed dispersal mutualisms has important implications for restoring understory herbaceous plant communities.

## Materials and Methods

### Study sites

We conducted this study across 20 paired sites in mesic hardwood forests in NAEDF. Each pair of sites included a remnant forest site (no history of clear cutting for agriculture ≥ 150 years) and a secondary site (regenerated from agricultural use (clear cut for pasture or plowing) 50-75 years). Remnant sites were located and verified using historic maps, aerial photographs, literature references, management reports, and land manager interviews (see details in Appendix S1). To mirror topographic conditions in the remnant sites, we selected secondary sites adjacent to or geographically close (within 32 km) to remnant sites. We grouped sites into three ecoregions (E1-E3) to account for regional variation (Appendix S1: Fig. S1).

At each study site, we set up three 50 m survey transects away from forest edges (> 100 m). Transects averaged 80 m from each other and were placed in areas with at least ∼75% deciduous tree cover. We created 5 m^2^ survey plots that alternated along transects with different plot types: seed removal, invertebrate community, and abiotic characteristic plots; vegetation plots; and ant habitat plots (Appendix S1: Fig. S2).

### Study system and species

*Aphaenogaster* are part of the *Aphaenogaster-rudis-fulva-texanus* species complex. (Umphrey 1996; Ellison et al. 2012; DeMarco & Cognato 2016). In our sites, we found *A. rudis* and *A. picea* (that are polyphyletic and challenging to delineate morphologically) and monophyletic *A. fulva* (Parker et al. 2021; Buono et al. 2022; Quartuccia & Buono, *unpublished data*) and refer to this group as *Aphaenogaster* sp. in our study. *Aphaenogaster* are abundant ants in NAEDF, with up to 2 nests per m^2^. They form single queen colonies (monogynous) with several hundred workers (Lubertazzi 2012). They are the primary dispersers of myrmecochores in NAEDF (Beattie & Culver 1981; Handel et al. 1981), including many species that we found at our sites, including *Trillium* sp., *Sanguinaria canadensis, Anemone acutiloba*, and *Asarum canadense (*Appendix S1: Table S3).

Other invertebrate organisms that are known to interact with myrmecochorous seeds in deciduous forests include other ant species (e.g., *Lasius americanus, Myrmica punctiventris*, and *Camponotus pennsylvanicus*) that are low-quality dispersers (Giladi 2006; Ness et al. 2009; Warren et al. 2015; Parker et al. 2021) (Appendix S1: Table S4). We also commonly observed the invasive slug belonging to the *Arion-subfuscus-fuscus* species complex interacting with seeds. This species complex is native to Europe and though both putative species are observed in North America, we refer to this slug as *A. subfuscus*, seeing as *A. subfuscus* is the more wide-ranging species (Pinceel et al. 2004). The slugs can be found in various habitats (forests, fields) and is a described pest, feeding on plants and fungi (Beyer & Saari 1978).

### Seed dispersal and invertebrate organisms interacting with seeds

We performed seed removal trials between late June to mid July 2018-2019 on non-rainy days to test if seed removal by ants differed between remnant and secondary sites. In each site, we set up 15 seed depots in seed removal/invertebrate community plots along three transects (Appendix S1: Fig. S2). Depots were located in the center of plots and consisted of a 10 cm^3^ petri dish placed on top of a paper index card (10×15 cm). Seeds from *A. canadense*, were collected when fruits were dehiscing from local sites. We chose *A. canadense*, as it is a common myrmecochore in this region (although not endemic to all sites; Appendix S1: Table S1) and is preferred by *Aphaenogaster* (Prior et al. 2015; Buono et al. 2022). To avoid unintentional introduction, seeds were frozen at -80 ºC for 24 hr. (to render seeds inviable) and then stored at 21 ºC. Seeds were thawed before placing them on depots and freezing is not known to impact ant interactions with seeds (Zelikova et al. 2008). On the day of the trials, we placed 8 seeds on depots. Wire mesh cages (7×12×12 cm) with 1.3 cm^2^ holes were secured over depots to exclude rodents but allow for invertebrate access. Depots were set out in the late morning (∼11:00 am) on non-rainy days. We observed depots 1/hr. for 3 hrs. During each observation, we recorded the number of seeds remaining, the presence and species of ants in depots, the type of interaction (carrying, handling), and the identification of any other invertebrates (i.e., slugs). Final observations were taken after 24 hrs., where we recorded the condition of the elaiosome (intact, partially removed, or completely scooped out), and the identification of any organisms on the depot (Meadley Dunphy et al. 2016; Parker et al. 2021).

We set out pitfall traps during the seed removal trials to compare *Aphaenogaster* sp. abundance, other ant abundance and diversity, and slug abundance between remnant and secondary sites. At each site we set up 30 pitfall traps (2 in each seed removal plot (Appendix S1: Fig S2)). The pitfall traps included a plastic cup (9 fluid oz., 7 cm tall, 9 cm diameter) filled with ∼ 3.5 oz of soapy water (with biodegradable soap) and depressed into the soil, top level with the ground. A 5 cm^2^ wire mesh grid with 1.3 cm^2^ openings covered the opening of the cups. Pitfall trap surveys were conducted on non-rainy days and left out for 24 hrs. Contents of traps were preserved in 70% ethanol for identification. We pinned specimens of ant species and identified all ants to the lowest taxonomic unit using regional keys (Ellison et al. 2012).

### Myrmecochore diversity and vegetation structure

We conducted vegetation surveys to compare differences in myrmecochore diversity and vegetation structure between remnant and secondary sites. Surveys were designed to compare forest composition and cover at several levels (understory, shrub, tree, and canopy) (Davison & Forman 1982). We set up four 1 m^2^ quadrats in the corners of each vegetative plot to measure herbaceous cover. We categorized plant species with seeds known to bear elaiosomes as myrmecochores and identified each to the lowest taxonomic unit (Handel et al. 1981; Bellemare et al. 2002) (Appendix S1: Table S3). In 5 m^2^ vegetative plots, we also measured shrub cover, tree basal area, percent canopy openness and identified tree species (see details in Appendix S1).

### Abiotic characteristics and potential ant habitat

In seed removal plots, we measured air temperature and three soil characteristics: soil pH, moisture, and temperature during the seed removal trials (see details in Appendix S1). We surveyed three known ant habitat types (leaf litter, decaying logs, and movable rocks (i.e., not large boulders)) in three plots (Appendix S1: Fig. S2) (see details in Appendix S1). For each habitat type, we recorded the presence of ant colonies (Appendix S1: Fig. S4).

### Statistical analysis

We performed binomial generalized linear mixed effects models (GLMM) on the number of seeds removed by ants and damaged by slugs (total elaiosome removed) per transect at the final observation time (24 hrs.). We included forest HLUC (remnant vs. secondary) and ecoregion as fixed effects and site nested within ecoregion as a random effect to account for spatial autocorrelation. We combined counts from two pitfalls per plot and performed a negative binomial GLMM (to account for overdispersion) on *Aphaenogaster* sp. abundance, other ant abundance, ant species richness. We estimated slug abundance as the combined number of slugs (or slug evidence) on seed depots and slug abundance in pitfall traps. We performed a negative binomial GLMM (to account for overdispersion) on slug abundance. Where ecoregion explained variation for main response variables (seed removal, organism abundance), we performed a Tukey’s post hoc test to compare differences among regions. When there was HLUC*ecoregion interactions, we compared remnant and secondary forests within ecoregions, performing a Bonferroni correction for multiple comparisons (Appendix S1: Table S2).

We performed linear mixed effects models (LMM) on percent myrmecochore cover, percent non-myrmecochore cover, shrub cover, tree basal area, and percent canopy openness at the transect level. We log-transformed (myrmecochore cover, shrub cover, and total basal area) to improve normality. We performed a negative binomial GLMM on species richness of myrmecochores at the transect level. Finally, we performed principal component analyses (PCA) on myrmecochore presence both at the site (PCA_S.myrmec_) and transect level (PCA_T.myrmec_) and performed LMMs on PC1_T.myrmec_ and PC2_T.myrmec_. Where ecoregion explained variation for myrmecochore richness and PC1_T.myrmec_, we performed a Tukey’s post hoc test to compare differences among regions. When there was HLUC*ecoregion interactions, we compared remnant and secondary forests within ecoregions, performing a Bonferroni correction for multiple comparisons (Appendix S1: Table S2).

We also performed LMMs on soil pH, moisture, temperature and on average air temperature at the transect level. We performed LMMs on rock surface area, leaf volume, and log volume, performing log-transformations on log volume and rock area.

To determine how organisms interacting with seeds and habitat factors influence observed variance in seed removal (See *Results*, Fig. 1), we performed path analyses. We constructed an *a priori* model with hypothesized pathways determined by known ecological interactions that included direct interactions between invertebrate organisms potentially interacting with seeds (*Aphaenogaster* sp., other ants, and slugs) and seed removal (Appendix S1: Fig. S5). We included direct interactions between “habitat” factors and the abundance of organisms interacting with seeds, and an indirect interaction between habitat and seed removal. We included direct interactions between other ants and *Aphaenogaster* sp. as other ants can affect seed dispersal if they interact antagonistically with *Aphaenogaster* sp. (Ness 2004; Warren et al. 2020). To maintain path analysis power, we created a composite habitat metric.

**Figure 1.**
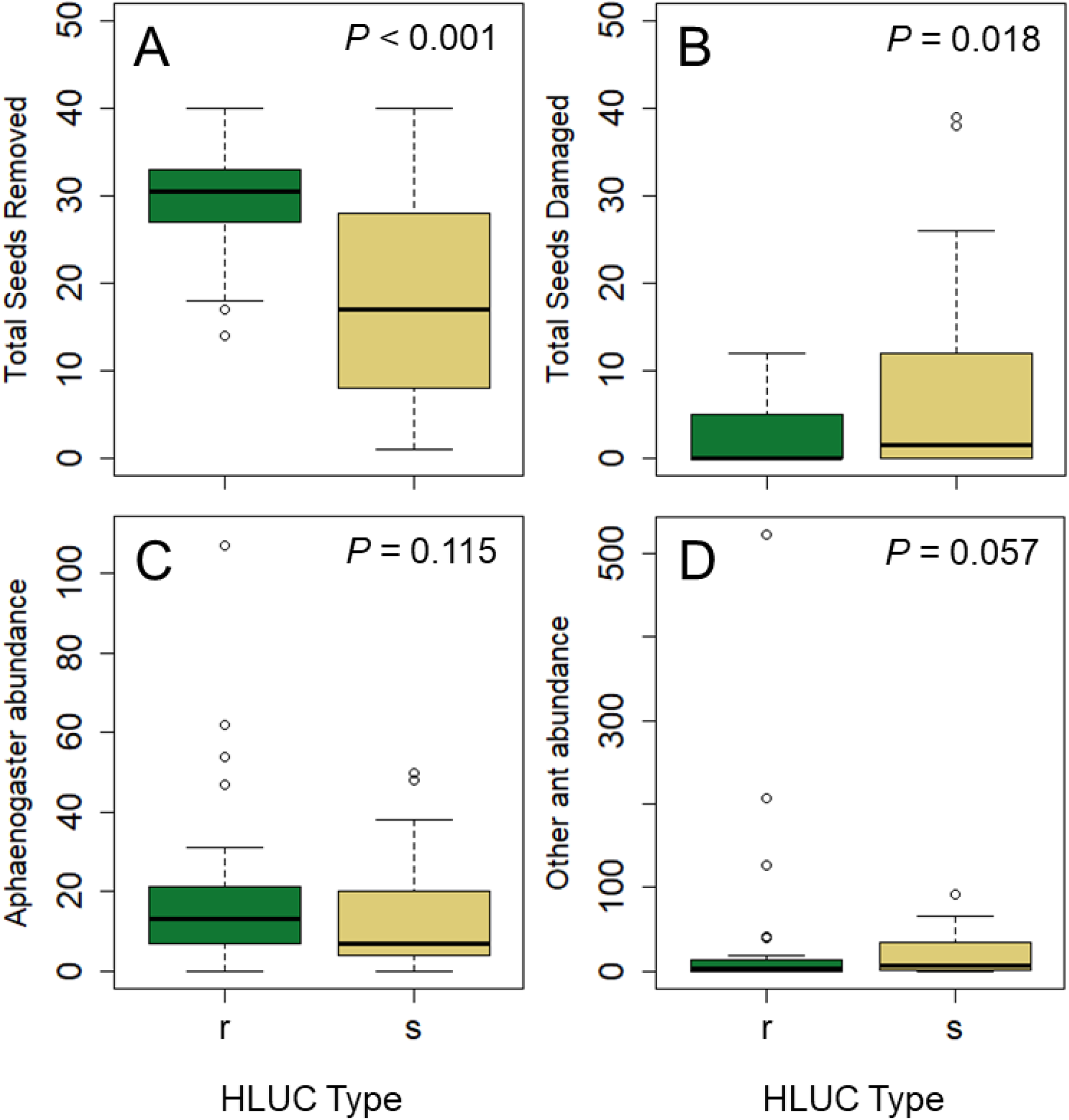
A) Seeds removed from depots, B) seeds with slug damage, C) *Aphaenogaster* sp. abundance, and D) other ant species abundance between remnant and secondary forests at the transect level (n = 60). Thick lines in box plots represent medians, boxes represent 1^st^ and 3^rd^ quartiles, whiskers represent minimums and maximums, and points represent outliers.

To create a composite habitat metric, we performed a correlation analysis among standardized variables representing forest structure, abiotic factors, ant habitat factors, organisms interacting with seeds (*Aphaenogaster* sp., other ant, and slug abundance), and seed dispersal (Appendix S1: Fig. S8). We performed a PCA on habitat factors with significant correlations to organisms and dispersal. PC_DM.hab_ was the first component and explained 35.4 percent of variation, and we used this component as the composite metric (Appendix S1: Fig. S10). Similarly, we ran seed dispersal path analyses separately for remnant forests and secondary forests and created composite habitat metrics for each (PC_DR.hab_ and PC_DS.hab_). We ran path analyses on myrmecochore cover for the combined dataset and separately for remnant and secondary forests (Appendix S1: Fig. S6). Myrmecochore cover was not significantly correlated with seed removal (see details in Appendix S1).

We used R for all analyses, including the following packages: MASS, lme4, stats, corrplot, lavaan, semplot, ggbiplot (Venables & Ripley 2002; Vu 2011; Rosseel 2012; Bates et al. 2015; Epskamp 2019; R Core Team 2021; Wei & Simko 2021).

## Results

### Seed removal and invertebrate organisms interacting with seeds

The number of seeds removed from depots was significantly higher in remnant forests compared to secondary forests (*P* < 0.001, Fig. 1A; see full statistical table, Appendix S1: Table S2). During the seed removal trials, the majority (> 90%) of ants removing seeds was *Aphaenogaster* sp. We also observed 9 ant species interacting with (but not removing) seeds (Appendix S1: Table S4). At 7 sites, we found *Nylanderia flavipes* (invasive ant) removing pieces of elaiosomes from seeds. The other main organism on depots was *A. subfuscus*, that fully scoops out elaiosomes, while not dispersing seeds (Meadley Dunphy et al. 2016).

In the pitfall traps, *Aphaenogaster* sp. abundance did not differ between forest HLUC type (*P* = 0.115; Fig. 1C). The abundance of other ant species also did not differ, but there was a trend for higher mean abundances in secondary forests (*P* = 0.057; Fig. 1D). We found 37 ant species, with no difference in richness between remnant and secondary forests (*P* = 0.783; Appendix S1: Table S2). Slug abundance was higher in secondary forests (*P* = 0.0025; Appendix S1: Table S2). We also found the number of seeds remaining with their elaiosome fully removed (due to slug damage) was higher in secondary forests (*P* = 0.0176; Fig.1B). We observed variation in seed removal, ant richness, and ant abundance among regions but no variation in slug abundance (Appendix S1: Table S2).

### Myrmecochore diversity and vegetation structure

We found 28 myrmecochore species (Appendix S1: Table S3), with total myrmecochore cover (*P* = 0.0316; Fig. 3A) and myrmecochore species richness being lower in secondary forests (*P* = 0.0041; Fig. 3B). In the site level myrmecochore PCA, we found PC1_S.myrmec_, accounted for 25.3% of variance with more forest indicator species at remnant sites, including *Dicentra* sp. (including *D. cucullaria*), *Tiarella cordifolia, Trillium* sp. (including *T. erectum*) along with nonsignificant indicator species *Anemone acutiloba* and *Claytonia caroliniana* (Appendix S1: Table S3). PC2_S.myrmec_ accounted for 16.1% of variance and was influenced by indicator species *T. grandiflorum, T. undulatum*, and *Uvularia sessilifolia* (Griffiths & McGee 2018) (Fig. 2). We found a difference in PC1_T.myrmec_ by forest HLUC (*P* = 0.0328; Appendix S1: Table S2), but PC2_T.myrmec_ did not differ between forest HLUC (Appendix S1: Fig. S3). Canopy openness was higher in secondary forests (*P* = 0.0084; Fig. 3C), but there were no differences in shrub cover, non-myrmecochore herbaceous understory cover, or tree basal area between forest types (Appendix S1: Table S2). There were ecoregion effects for myrmecochores, with myrmecochore richness being higher in E3 than in E1 and E2 (Fig 3B; Appendix S1: Table S2).

**Figure 2.**
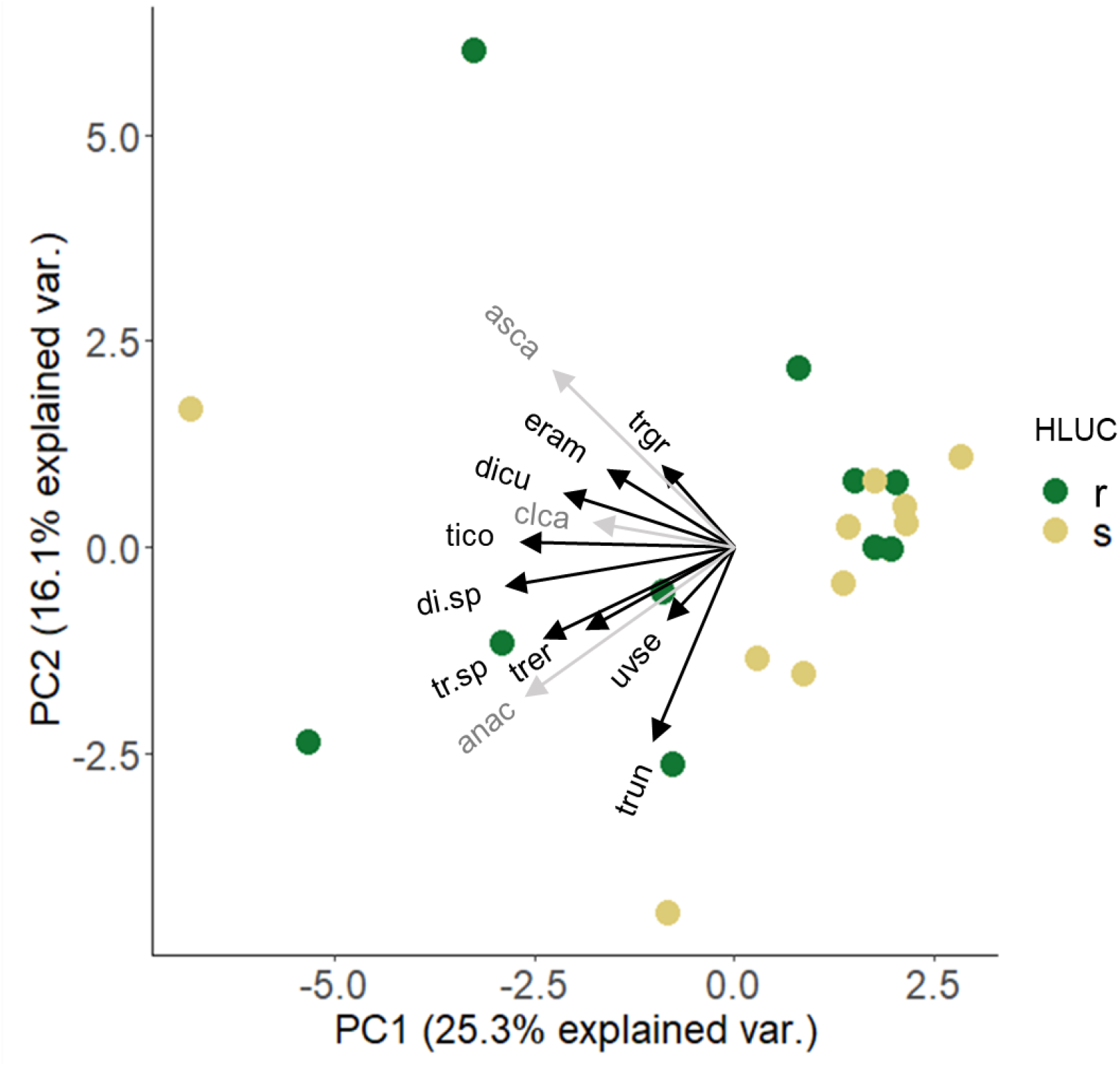
Myrmecochore species site level biplot of principal component analysis (PCA). Remnant transects (r) represented by green symbols and secondary (s) are tan symbols. Remnant sites are more diverse (spread out) compared to the majority of secondary sites clustered on the right side of PC1_S.myrmec_. PC1_S.myrmec_ explains 25.3% of variance and PC2_S.myrmec_ 16.1%. Black arrows represent indicator species and gray arrows nonsignificant indicator species (Griffiths & McGee 2018). Species acronyms found in Appendix S1: Table S3.

**Figure 3.**
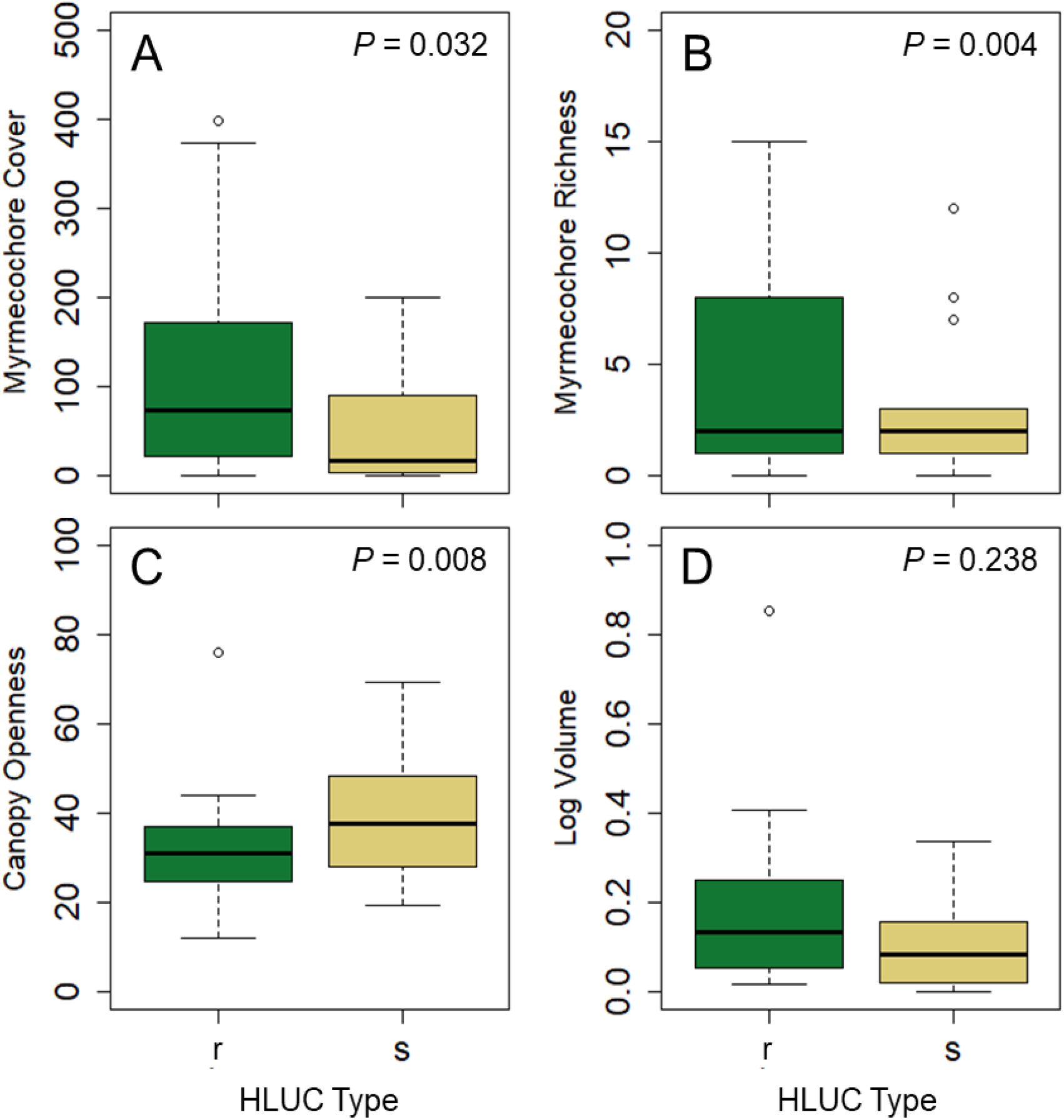
A) Myrmecochore species cover, B) myrmecochore species richness, C) canopy openness, D) log habitat volume at the transect level (n = 60) in remnant (r) and secondary (s) forests. Thick lines in box plots represent medians, boxes represent 1st and 3rd quartiles, whiskers represent minimums and maximums, and points represent outliers.

### Abiotic characteristics and potential ant habitat

We did not find differences in average soil temperature, soil pH, soil moisture, or air temperature between HLUC (Appendix S1: Table S2). We also found no differences in leaf litter volume, log volume, or rock surface area between HLUC (Fig. 3D). Across both types of HLUC, we found a slight difference in habitat types occupied by ant colonies, preferring log over leaf litter and rock (Appendix S1: Fig. S4).

### Path analyses and correlation analyses

In the combined seed dispersal path model, we found significant and positive interactions between *Aphaenogaster* sp. abundance and other ant abundance (0.72 ± 0.16; *P* < 0.001; Fig. 4A). *Aphaenogaster* sp. abundance had a significant positive effect on seed removal (0.47 ± 0.16; *P* < 0.01) and slug abundance had a significant negative effect (−0.36 ± 0.11; *P* < 0.01). The abundance of other ants had no effect on seed removal (Fig. 4A). PC_DM.hab_ had a significant negative effect on the abundance of other ants (0.38 ± 0.12; *P* < 0.01; Fig. 4A). Given the factors that contributed to PC_DM.hab_, ant abundance is higher at sites with relatively lower soil moisture, leaf litter, and air temperature, and higher soil pH, soil temperature, and herbaceous cover (Fig. 4A; Appendix S1: Fig S8). The final model was supported by the data (χ^2^ = 1.40; d.f. = 2; *P* = 0.49). A *P* value > 0.05 suggests the data are consistent with the hypothesized model.

**Figure 4.**
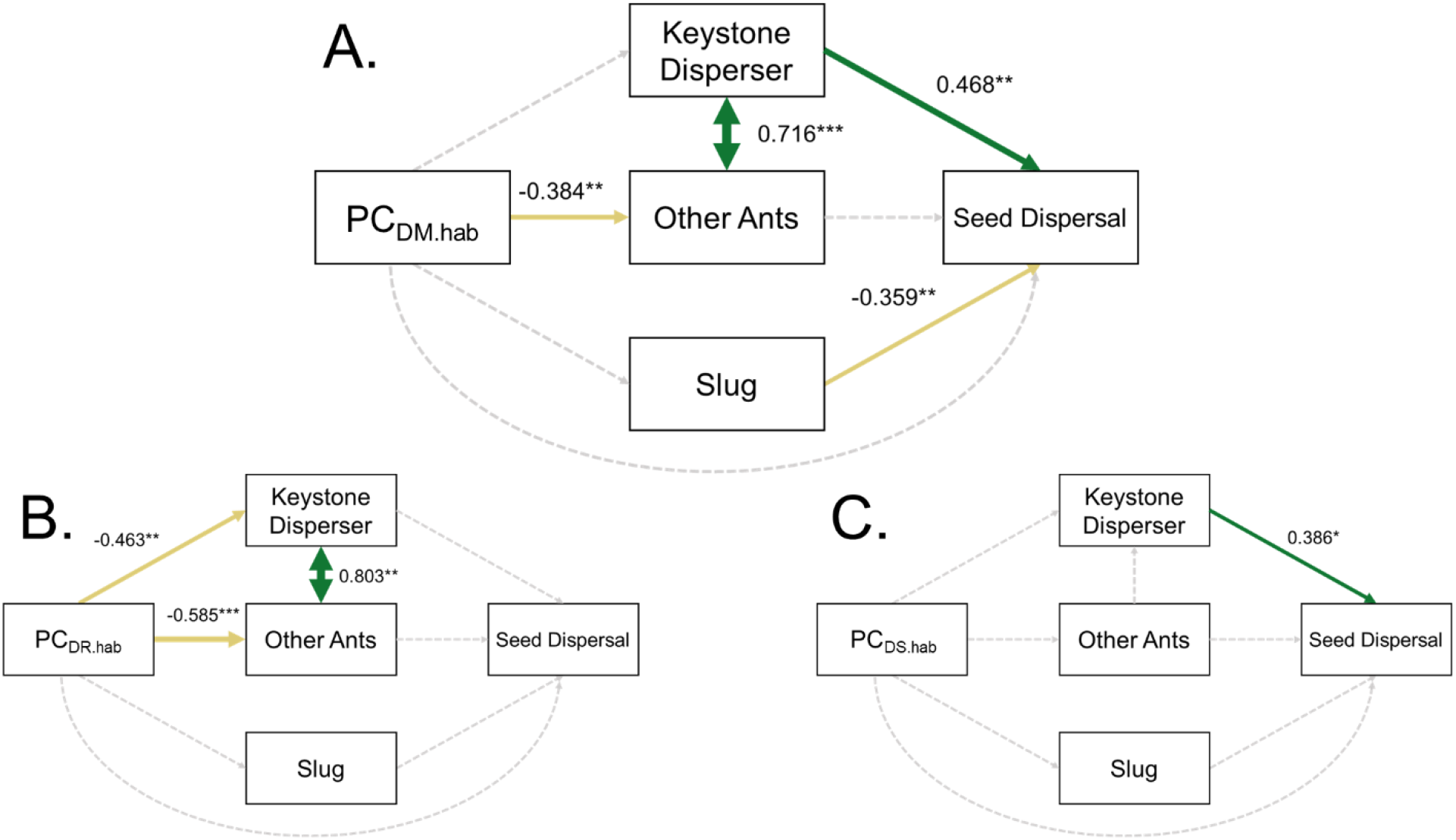
Seed dispersal path diagram with standardized path coefficients reported next to the arrows for A) all forests (combined) B) remnant forests and C) secondary forests. Green and tan solid arrows indicate significant positive and negative pathways respectively. Thickness of arrows are proportional to the standardized path coefficient’s strength. Non-significant pathways with path coefficients less than 0.1 are given in dashed gray lines. Significant differences represented by symbols (* > 0.05, ** > 0.01, *** > 0.001).

The remnant forest path model also revealed that *Aphaenogaster* sp. abundance and other ant abundance had a significant, positive interaction (0.80 ± 0.23; *P* < 0.01; Fig. 4B). However, in this case, the abundance of organisms had no significant direct effect on seed removal. In the remnant path, PC_DR.hab_ had significant effects on *Aphaenogaster* sp. and other ants (0.46 ± 0.16, *P* < 0.01; 0.59 ± 0.15, *P* < 0.001; Fig. 4B), again with soil conditions (lower moisture and higher temperature) influencing ant abundance (Appendix S1: Fig. S8). The secondary forest path model revealed that *Aphaenogaster* sp. abundance had a direct positive effect on seed removal (0.39 ± 0.15; *P* < 0.05; Fig. 4C), but no other interactions were significant. Secondary PC_DS.hab_ had no effect on organism abundance (Fig. 4C; Appendix S1: Fig. S8). The final models were both supported by the data (χ ^2^ = 1.01, d.f. = 2, *P* = 0.60; χ ^2^ = 1.98, d.f. = 2, *P* = 0.37). The combined and separate path analyses showed weak effects of organisms’ abundance and habitat factor impacts directly on myrmecochore cover (Appendix S1: Fig S6). However, our correlation analysis shows significant positive effect of soil pH and other herbaceous (non-myrmecochore) cover on myrmecochore cover in the combined and remnant forests, but not the secondary forests (Appendix S1: Fig. S9).

## Discussion

Our work shows that seed dispersal by ants, a vital ecosystem function in NAEDF ecosystems, is partially resilient to disturbances from HLUC with variation in recovery trajectories. Secondary forests had lower and more varied rates of seed dispersal than remnant forests, resulting from altered interactions with mutualists and antagonists. High-quality mutualists, *Aphaenogaster* sp., were the primary dispersers of seeds and variation in their abundance contributed to variation in seed dispersal. Other ant species did not affect seed dispersal, but invasive slugs were more abundant and slug-induced seed damage more prevalent in secondary forests, with a negative relationship between slug abundance and seed dispersal. Interestingly, variation in *Aphaenogaster* sp. abundance did not influence seed dispersal in remnant forests (only in secondary forests), suggesting that seed dispersal function is stable in remnant forests, but variable in secondary forests, where dispersal quality is dependent on *Aphaenogaster* sp. abundance. Myrmecochore cover and richness was lower in secondary forests, but not influenced by *Aphaenogaster* sp. abundance. Taken together, secondary forests vary in functional resilience, where dispersal is reduced in some forests with low mutualist abundance, but intact where mutualists are abundant. While variation in functional resilience may not be a direct mechanism of low recovery of myrmecochory diversity in secondary forests, animal-mediated seed dispersal, including myrmecochory, is known to be important for plant fitness, distribution, and community structure (Kalisz et al. 1999; Canner et al. 2012; Prior et al. 2015). As result, functional resilience of seed dispersal interactions should be considered when restoring myrmecochore communities.

We predicted that seed dispersal would be lower in secondary forests due to lower abundances of mutualistic partners, *Aphaenogaster*. While we found lower rates of seed dispersal in secondary forests, *Aphaenogaster* sp. was present in all forests, with no difference in abundance between HLUC. This suggests that *Aphaenogaster* are somewhat resilient to widespread forest clearing. Similarly, previous work found no difference in abundances of *Aphaenogaster* between forests of differing HLUC (Mitchell et al. 2002; Kiel et al. 2020). Despite finding no difference, variation in *Aphaenogaster* sp. abundance was the primary determinant of seed dispersal variation in all forests. Interestingly, the sensitivity of seed dispersal to mutualist abundance differed between remnant and secondary forests. In remnant forests, seed dispersal variation was not influenced by *Aphaenogaster* sp. abundance, but dispersal in secondary forests was sensitive to changes in *Aphaenogaster* sp. abundance. These different relationships suggest that seed dispersal function is stable and intact in remnant forests, but variable in secondary forests, with high dispersal occurring in secondary forest patches with high *Aphaenogaster* sp. abundance.

In our combined and remnant seed dispersal path analysis, we found an effect of habitat conditions on other ant and *Aphaenogaster* sp. abundance. Our findings support previous studies that show that *Aphaenogaster* distribution is determined by microhabitat conditions, specifically that they do not occupy soil with high moisture (Warren et al. 2010; 2011; 2012). In secondary forests, *Aphaenogaster* sp. abundance was not influenced by habitat variation. This could be due to greater microhabitat homogenization, a legacy of widespread forest clearing (Flinn & Marks 2007). Additionally, variation in nest disturbance or recolonization ability from local source populations may be more important in influencing abundance than microhabitat conditions.

We also predicted variation in antagonists would influence seed dispersal and found higher abundances of *A. subfuscus*, and higher rates of slug-caused seed damage in secondary forests. In the combined path analysis, we found a negative relationship between slugs and seed dispersal. Elaiosome removal by slugs decreases dispersal by ants by removing the attractive food reward and disrupting the interaction (Meadley Dunphy et al. 2016). Slugs might be more abundant in secondary forests if they have increased access through forest fragmentation and proximity to other habitats, like old fields (Beyer & Saari 1978; Kozłowski 2009). In a previous study, we found higher slug abundances and elaiosome damage at forest edges than in forest interiors (Parker et al. 2021). Slug abundance might also be higher in secondary forests due to changes in environmental conditions, but we found few consistent differences in microhabitat conditions between HULC. We found no effect of habitat factors on slug abundances in our path analyses, but soil temperature had a positive correlation with slug abundance, and there was a negative relationship with soil moisture (Appendix S1: Fig. S8). Future studies investigating the mechanisms leading to increased slug presence in secondary forest would be useful, especially when considering restoring understory plants reliant on seed dispersal mutualisms.

In all of our path analyses, other ant species (species other than *Aphaenogaster* sp.) abundance did not have a direct effect on seed dispersal but had a strong positive interaction with *Aphaenogaster* sp. abundance. This suggests that overall, other ant species do not have a direct antagonistic interaction with seeds or mutualist partners. This is an interesting finding, as it is generally predicted that other ants negatively affect dispersal directly (by being antagonistic or low quality partners) or indirectly by outcompeting the good disperser (Ness 2004; Giladi 2006; Ness et al. 2009; Prior et al. 2020; Parker et al. 2021). Habitat factors in both the combined and remnant path analysis influenced other ant species abundance, which suggests that the other ant species were more abundant in microhabitats that also favored *Aphaenogaster* sp.. During the seed removal trials, we only observed two other ant species interacting with, but not removing, seeds: the native species *Lasius americanus* and the invasive species *Nylanderia flavipes*, with the latter occurring at high abundances and removing parts of elaiosomes. Taken together, other ants do not seem to be largely antagonistic, but also do not always contribute to seed dispersal function.

Previous work examining HLUC on myrmecochory in NAEDF found that patch size and historical land use intensity influences ant community abundance and composition (Mitchell et al. 2002). While this previous study found little effect of HLUC on *Aphaenogaster* abundance, they did not quantify how the abundance or presence of ants directly affected seed dispersal function. More recent work directly assessing seed removal by ants in forests that differed in HLUC demonstrated that rates of removal did not differ (Kiel et al. 2020). While this work contributes to our understanding of HLUC effects on ant-mediated seed dispersal, it was limited in spatial scope, not accounting for variation at the landscape level, with only three forests in a narrow portion of the range of this mutualism. Here, we covered a larger portion of this mutualism’s range and in doing so demonstrated variation in functional resilience at the landscape level. We also found ecoregion effects on ant abundances, forest structure, and abiotic factors which could contribute to variation in functional resilience across space. To this end our study does not contradict either previous study, rather our increased scale and direct measurement of seed dispersal function expands the scope, revealing variation in functional resilience despite *Aphaenogaster* being present in secondary forests.

We found that total myrmecochore cover and richness were lower in secondary sites. This finding is similar to previous work that finds lower abundances of myrmecochore species in forests that have been previously cleared (Bellemare et al. 2002; Mitchell et al. 2002; Vellend 2005; Griffiths & McGee 2018). In our myrmecochore PCA, myrmecochore species composition differed between secondary and remnant forests. Remnant forests had more cover and species richness, and particularly higher presence and abundance of specific remnant forest indicator species (Griffiths & McGee 2018). We found that myrmecochore cover was not influenced by *Aphaenogaster* sp. abundance in our combined path analysis and that myrmecochore cover and seed dispersal is not significantly correlated. This suggests that factors other than ant mutualists and seed dispersal are contributing to variation in myrmecochore cover. Habitat factors such as soil pH, nutrient availability, and organic matter content are all known to be altered by forest clearing and agricultural disturbances and could be contributing to lower myrmecochore cover in secondary forests (Koerner et al. 1997; Dyer 2010). However, we found no difference in soil characteristics between forests with different HLUC which suggests soil resilience in some disturbed forests. We found secondary forests had higher canopy openness, which could impact other abiotic factors that contribute to myrmecochore cover such as light availability. Other studies suggest that low myrmecochore richness and cover may also be affected by recruitment limitation from source populations (Bellemare et al. 2002; Flinn & Vellend 2005).

One consideration in our study is that forests varied in myrmecochore cover, which could affect seed dispersal. For example, *Aphaenogaster* colonies are known to become satiated with myrmecochorous seeds (Heithaus et al. 2005), and forests with more myrmecochores could mean that ants forage less for seeds. However, we found no relationship between myrmecochore cover and seed dispersal. Another source of variation we did not account for, is that we pooled *Aphaenogaster* species given that they are challenging to tell apart in the field. Emerging work, including our own, suggests that mutualistic partner identity (among *Aphaenogaster* putative species and populations) can affect seed dispersal function (Warren & Bradford 2014; Prior et al. 2015; Meadley Dunphy et al. 2020; Prior et al. 2020; Buono et al. 2022). Also, we controlled for rodent impacts on seed dispersal, but variation in this antagonistic interaction could also be contributing to how HLUC affects this mutualism (Ness & Morin 2008).

Variation in legacy effects and resilience can cause variable recovery trajectories, making predicting resiliency or trying to reverse impacts of disturbances difficult and complex (Suding et al. 2004). Specifically for ant-mediated seed dispersal in NAEDF, while mutualist ant partners are present in secondary forests, suggesting some level of resilience, there is variation in their abundance and function. Other studies show variation in the resiliency of diversity post disturbance (Steadman 1997; Elmqvist et al. 2002; Sabatini et al. 2014), but less is known about the resiliency of functionally important interactions that diversity relies on (Oliver et al. 2015, (Mitchell et al. 2002; Kaiser-Bunbury et al. 2017; García et al. 2018). Uncovering how interactions that species rely on are resilient to disturbance is critical to understand mechanisms of slow recovery or predict if functions will be intact for proposed active restoration.

Our research provides implications for restoration efforts. First, we emphasize the importance of preserving remaining remnant forest ecosystems to provide critical source populations for recovery. Second, given that not all secondary forests are resilient to historical forest clearing suggests that forest patches with intact seed dispersal interactions might be prioritized for active restoration of understory plants, or there may need to be efforts to augment or enhance this interaction in some forests. While the presence of seed dispersal function and mutualistic ants do not directly determine plant community resilience, their documented importance on understory plant populations and communities means that maintenance of this function will be essential to conserving understory plant communities.

## Acknowledgements

We thank Shannon Meadley Dunphy, Wyatt J. Parker, Dudley Britton, and Alfonso J. Buono for help in the field and Melina Brunelli, Lena Eder, Roni Friedman, and Kassidy Robinson for help in the lab. We thank the following land managers: Rutgers, The State University of New Jersey; U.S. National Park Service; The Nature Conservancy; Pennsylvania Department of Conservation and Natural Resources; Pennsylvania Game Commission; Cornell Botanical Gardens; New York Botanic Garden Thain Family Forest; and New York City Department of Parks & Recreation. Funding came from Binghamton University (BU), BU Center for Integrated Watershed Studies (CMB), BU Provost’s Scholarship (CMB) and the Torrey Botanical Society (CMB). This work was conducted on Haudenosaunee, Susquehannock, Lenni-Lenape, and Munsee Lenape ancestral lands; land that Indigenous communities were forcibly removed from. We acknowledge Indigenous peoples as the original stewards of this land and honor the lasting relationship between these nations and the land. We offer this land acknowledgement to affirm Indigenous sovereignty, history, and experiences connected with these regions, and we hope to advocate for increased Indigenous recognition within the scientific and research community.

## Author contributions

CMB and KMP conceived the idea and designed the research; CMB, KMP, CG, and JS conducted the fieldwork; CMB, JF, WS, CG, and JS performed lab-work; CMB analyzed the data with input from KMP; CMB wrote the manuscript with input from KMP.

