## Appendix S1 for "Historical forest disturbance results in variation in functional resilience of seed dispersal mutualisms"

**Historical forest disturbance results in variation in functional resilience of seed dispersal mutualisms**

Carmela M. Buono, Jesse Lofaso, Will Smisko, Carly Gerth, John Santare, Kirsten M. Prior

Ecology

**Supplementary Methods and Results**

*Study sites*

We conducted this study across 10 paired sites in mesic hardwood forests in the northeastern mid-Atlantic region of North America. Each pair of sites included a remnant forest site (a forest with no history of clear cutting for agriculture within the past 150 or more years) and a secondary site (a forest regenerated from agricultural use (clear cut for pasture or plowing) within the past 50-75 years). Remnant sites were located and verified using historic maps (1990s), aerial photographs (1940s), literature references, management reports, and land manager interviews. Secondary sites were selected to mirror topographic conditions in the remnant sites by choosing sites adjacent to remnant sites when possible or geographically close (within 32 km). We grouped sites into three ecoregions (E1-E3) to account for regional variation (Appendix S1: Fig. S1). Ecoregions (Table S1; Fig. S1) were determined by Level III Environmental Protection Agency ecoregions and by comparing dominant tree species (Fig. S7). Sites varied in tree canopy, with dominant trees including *Acer rubrum*, *Tsuga canadensis*, *A. saccharum*, *Quercus rubra*, and *Fagus grandifolia* (Appendix S1: Table S1).

At each study site, we set up three 50 m survey transects away from forest edges (> 100m from edge). Transects averaged 80 m from each other and were placed in areas with at least ~75% deciduous tree cover. We created 5 $m^2$ survey plots that alternated along transects (to prevent trampling) in plot types: seed removal, invertebrate community, and abiotic characteristic plots; vegetation plots; and ant habitat plots (Appendix S1: Fig. S2).

*Myrmecochore diversity and vegetation structure*

We conducted vegetation surveys in plots along transects to compare differences in myrmecochore diversity and vegetation structure between remnant and secondary study sites. Surveys were performed from late May to early June from 2018-2019, which allowed us to observe evidence of myrmecochore species that emerged earlier as well as the majority of the growing season.

Vegetation surveys were performed along the 3 transects at each site, in 5$m^2$ plots on alternating sides of the transect line that did not overlap with seed dispersal plots (Fig. S2). We designed the vegetation surveys to compare forest composition and cover at several levels (understory, shrubs, trees, and canopy) and were developed from other long-term studies (Davison & Forman 1982). To measure percent canopy openness, a spherical concave mirror densiometer was used in the center of each vegetation survey plot. All trees within the canopy layer directly above the plot were identified to species. To measure the trees at each site, every tree > 3 cm diameter within the 5 $m^2$ plot was measured for diameter at breast height (DBH) and identified to species (Table S1). Total basal area for each 5 $m^2$ plot was calculated using the

DBH data. To measure the shrub layer, we used a line intercept to record relative cover of any woody plant between 0.3-3 meters tall. We ran a 10 m transect originating from the transect line and overlapping with the 5 $m^2$ vegetation survey plot and recorded any shrub that intersected the 10 m line. We recorded the length that each shrub covered the transect (by the shape of the entire plant, including leaves, stems, and spaces between).

We set up four 1 $m^2$ quadrats in the corners of each vegetative plot to measure the understory layer. We measured herbaceous cover (graminoids, forbs, ferns, and fern allies) within the quadrats. We categorized any plant species with seeds known to bear elaiosomes as myrmecochores and identified each species to the lowest taxonomic unit (Handel et al. 1981; Lengyel et al. 2009) (Table S1). Additionally, we recorded the presence of any myrmecochores species found in the 5 $m^2$ plot but not recorded in the 1 $m^2$ quadrats.

*Abiotic characteristics and potential ant habitat*

We measured three soil characteristics: soil pH, moisture, and soil temperature in the seed removal trail plots during the seed removal trials. Each characteristic was measured 5 times in the middle of the plot and then averaged. We measured soil temperature with a soil thermometer placed 10 cm into the soil. We measured soil moisture and soil pH with a soil acidity and moisture meter (Kelway HB-2 Soil Acidity and Moisture Tester, Pacific Star Corporation, Texas). During the seed removal trial, we measured ground air temperature with iButton temperature loggers (Thermochron, On Solution, Sydney, Australia) hung ~30 cm off the ground in a mesh bag that were not placed in direct sunlight. Temperature readings were taken every 60 minutes for the 24 hours.

We conducted potential ant habitat surveys in three 5 $m^2$ plots along transects that did not overlap with the vegetation or seed removal trials, from June-July 2019-2020 (Fig. S2). Within each plot, we set up 2 1x5$m^2$ belt transects. We quantified ant habitat types in the belt transects by measuring the volume of leaf litter piles (length x width x height) and the cylindrical volume of logs (3.14 x width x 2 x height). In the same belt transects, we measured total surface area of movable rock habitat (i.e., not large boulders) by estimating percent cover in a 1x1$m^2$ quadrat. We focused on these three ant habitat types because they are known to be preferred habitat for the keystone dispensers *Aphaenogaster* sp. (Lubertazzi 2012). For each ant habitat type, we also indicated the presence of ant colonies. For logs, we quantified the level of decay by measuring the distance (cm) a metal skewer (0.9 cm diameter) could be punctured into the center of the log with moderate force.

*Statistical analysis*

We created composite habitat metrics by performing a correlation analysis among standardized variables representing forest structure, abiotic factors, ant habitat factors, organisms interacting with seeds (*Aphaenogaster* sp. abundance, other ant abundance, and slug abundance), and seed dispersal (Appendix S1: Fig. S8). We then selected all habitat-related factors that had a significant correlation with any of the organism and seed dispersal response variables (Appendix S1: Fig. S8). We performed a PCA on these select habitat-related factors to create a composite habitat metric ($PC_{DM.hab}$) (that we put in the path analysis) that represented habitat factors that influence organism abundance and dispersal. Six variables were included in the $PCA_{DM.hab}$ (Appendix S1: Fig. S10). $PC_{DM.hab}$ was the first component and explained 35.4 percent of variation. Positive values represent high soil moisture, leaf volume, and air temperature, and low soil pH, soil temperature, and other (non-myrmecochore) herbaceous cover. Similarly, we ran path analyses separately for remnant forests and secondary forests and created composite habitat metrics for each ($PC_{DR.hab}$ and $PC_{DS.hab}$).

We also ran path analyses on myrmecochore cover for the combined dataset and separately for remnant and secondary forests (Appendix S1: Fig. S6) using similar paths as described above. Similarly, we created composite habitat metrics, this time including habitat factors that had a significant relationship with myrmecochore cover and organism abundance. The composite habitat metric for the combined myrmecochore path analysis shares the same name as the combined seed dispersal path analysis ($PC_{DM.hab}$) as it had the same group of significant habitat factors (Appendix S1: Fig. S9, S10). Myrmecochore cover was not significantly correlated with seed dispersal.

**Table S1.** Study site details including land managers, site code, ecoregion (E1-E3, see Fig. S1) latitude, longitude, elevation, soil profile, previous land history resources, dominant tree species, and if *Asarum canadense* (asca) is present at the site.

| Site code | Site manager | ER | Latitude, Longitude | Elevation (m) | Soil profile | Land use history sources | Dominant Tree | asca present |
|---|---|---|---|---|---|---|---|---|
| 1R | Rutgers, State University of New Jersey | ER1 | 40.499077, -74.561999 | 22 | Fine, mixed | Historical maps; Davis 2003; Kershner & Leverett 2004; and manager verification | *Quercus alba, Acer rubrum, Fagus grandifolia* | |
| 1S | Rutgers, State University of New Jersey | ER1 | 40.487215, -74.569072 | 25 | Loamy-skeletal | Historical maps and manager verification | *Quercus rubra* | |
| 2R | U.S. National Park Service | ER1 | 40.765317, -74.523400 | 198 | Fine-loamy/ Loamy-skeletal | Historical maps and Kershner & Leverett 2004 | *Quercus alba* | |
| 2S | U.S. National Park Service | ER1 | 40.765965, -74.540367 | 177 | Fine-loamy | Historical maps | *Quercus velutina* | |
| 3R | The Nature Conservancy | ER3 | 41.755278, -75.892778 | 477 | Coarse-loamy/ Loamy-skeletal | Historical aerial photographs; Davis 2003; Kershner & Leverett 2004; and manager verification | *Acer rubrum, Tsuga canadensis* | yes |
| 3S | The Nature Conservancy | ER3 | 41.881854, -75.504037 | 537 | Coarse-loamy/ Loamy-skeletal | Historical aerial photographs and manager verification | *Acer saccharum, Prunus serotina* | yes |
| 4R | Pennsylvania Dept. of Conservation and Natural Resources | ER3 | 41.909733, -75.863867 | 388 | Coarse-loamy/ Loamy-skeletal | Historical aerial photographs; Davis 2003; Kershner & Leverett 2004; and manager verification | *Tsuga canadensis, Fagus grandifolia* | |
| 4S | Pennsylvania Game Commission | ER3 | 41.944174, -75.709691 | 334 | Coarse-loamy | Historical aerial photographs and manager verification | *Acer rubrum, Quercus rubra* | |
| 5R | Cornell Botanical Gardens | ER3 | 42.421484, -76.324251 | 467 | Fine-loamy | Historical aerial photographs and university documents | *Tsuga canadensis, Acer saccharum* | |
| 5S | Cornell Botanical Gardens | ER3 | 42.353389, -76.381000 | 482 | Fine-loamy | Historical aerial photographs and university documents | *Acer rubrum* | |
| 6R | Cornell Botanical Gardens | ER3 | 42.333257, -76.664323 | 454 | Coarse-loamy | Historical aerial photographs and university documents | *Tsuga canadensis, Fagus grandifolia* | |

| | | | | | | | | |
|---|---|---|---|---|---|---|---|---|
| 6S | Cornell Botanical Gardens | ER3 | 42.340397, -76.662643 | 444 | Fine-loamy | Historical aerial photographs and university documents | *Acer rubrum, Fraxinus sp.* | |
| 7R | Cornell Botanical Gardens | ER3 | 42.455004, -76.450653 | 263 | Fine-loamy | Historical aerial photographs; Davis 2003; and university documents | *Acer saccharum, Platanus occidentalis* | yes |
| 7S | Cornell Botanical Gardens | ER3 | 42.458578, -76.447877 | 305 | Fine-loamy | Historical aerial photographs and university documents | *Fagus grandifolia, Carya glabra, Tsuga canadensis* | yes |
| 8R | New York Botanic Garden Thain Family Forest | ER2 | 40.864050, -73.876251 | 34 | Coarse-loamy | Davis 2003 and manager verification | *Tsuga canadensis, Quercus rubra* | |
| 8S | New York City Department of Parks & Recreation | ER2 | 40.906215, -73.892388 | 27 | Coarse-loamy | Manager verification | *Quercus rubra, Liriodendron tulipifera* | yes |
| 9R | Pennsylvania Department of Conservation and Natural Resources | ER2 | 41.298667, -76.267017 | 391 | Coarse-loamy | Historical aerial photographs; Davis 2003; Kershner & Leverett 2004; and manager verification | *Betula lenta, Nyssa sylvatica* | |
| 9S | Pennsylvania Department of Conservation and Natural Resources | ER2 | 41.334833, -76.265717 | 671 | Coarse-loamy | Historical aerial photographs and manager verification | *Acer rubrum, Fagus grandifolia, Prunus serotina* | yes |
| 10R | Pennsylvania Department of Conservation and Natural Resources | ER3 | 40.781134, -75.292735 | 136 | Fine-loamy | Historical aerial photographs; Davis 2003; Kershner & Leverett 2004; and manager verification | *Liriodendron tulipifera, Acer rubrum* | yes |
| 10S | Pennsylvania Department of Conservation and Natural Resources | ER3 | 40.789112, -75.290664 | 159 | Loamy-skeletal | Historical aerial photographs and manager verification | *Fraxinus sp., Acer rubrum, Carpinus caroliniana* | |

*Soil type and profile data were collected using U.S.G.S. Web Soil Survey.

**Table S2** Statistics for response variables, including d.f. and *P* values. Linear mixed effects models (LMM) and generalized linear mixed effects models (GLMM) were performed with historical land use change (HLUC) and ecoregion as fixed factors and site nested in ecoregion as a random effect. Bolded values represent significant effects. F values are reported for LM and Deviance and Residual Deviance for negative binomial GLMs. Tukey's posthoc tests were performed among ecoregions when ecoregion was significant for main response variables (*). When HLUC*Ecoregion was significant, we compared HLUC within each region using Bonferroni corrections to determine significance (α = 0.017). (E1-E3, see Fig. S1; Table S1), r = remnant, s = secondary, ns = non-significant).

| **Response variable** | **Factors** | **d.f.** | **F value or Deviance (Residual Deviance)** | ***P*-value** | **Direction of difference, and within region comparison (α = 0.017)** |
|---|---|---|---|---|---|
| ***Seed removal** | **HLUC** | **1, 54** | **25.47** | **< 0.001** | **r > s** |
| | **Ecoregion** | **2, 54** | **10.67** | **< 0.001** | **E1, E2 > E2, E3** |
| | **HLUC*Ecoregion** | **2, 54** | **7.15** | **0.002** | E1 (ns)<br>E2 (ns)<br>**E3 (s > r) (< 0.001)** |
| ***Seed slug damage** | **HLUC** | **1, 54** | **6.00** | **0.018** | **r < s** |
| | Ecoregion | 2, 54 | 1.23 | 0.300 | |
| | HLUC*Ecoregion | 2, 54 | 1.41 | 0.253 | |
| ***Aphaenogaster* abundance** | HLUC | 1, 54 | 2.48 (85.28) | 0.115 | |
| | **Ecoregion** | **2, 54** | **17.79 (67.49)** | **< 0.001** | **E1, E2 > E1, E3** |
| | HLUC*Ecoregion | 2, 54 | 0.25 (67.24) | 0.884 | |
| ***Other ant abundance** | HLUC | 1, 54 | 3.62 (1445.23) | 0.057 | |
| | **Ecoregion** | **2, 54** | **67.31 (77.92)** | **< 0.001** | **E1, E2 > E1, E3** |
| | **HLUC*Ecoregion** | **2, 54** | **10.37 (67.55)** | **0.006** | E1 (ns)<br>E2 (ns)<br>E3 (ns) |
| ***Ant richness** | HLUC | 1, 54 | 0.08 (122.35) | 0.783 | |
| | **Ecoregion** | **2, 54** | **63.05 (59.30)** | **< 0.001** | **E1, E2 > E3** |
| | HLUC*Ecoregion | 2, 54 | 3.07 (56.23) | 0.216 | |
| ***Slug abundance** | **HLUC** | **1, 54** | **9.14 (55.40)** | **0.003** | **r < s** |
| | Ecoregion | **2, 54** | 2.07 (53.33) | 0.355 | |
| | HLUC*Ecoregion | 2, 54 | 4.35 (48.98) | 0.114 | |
| ***Myrmecochore cover** | **HLUC** | **1, 54** | **4.87** | **0.032** | **r > s** |
| | Ecoregion | 2, 54 | 1.15 | 0.326 | |
| | HLUC*Ecoregion | 2, 54 | 0.63 | 0.534 | |
| ***Myrmecochore richness** | **HLUC** | **1, 54** | **8.24 (93.76)** | **0.004** | **r > s** |
| | **Ecoregion** | **2, 54** | **35.85 (57.91)** | **< 0.001** | **E1, E2 < E3** |
| | HLUC*Ecoregion | 2, 54 | 2.67 (55.24) | 0.263 | |
| ***PC1**$_{T.myrmec}$ | **HLUC** | **1, 54** | **4.80** | **0.033** | r > s |
| | **Ecoregion** | **2, 54** | **3.28** | **0.045** | ns |
| | HLUC*Ecoregion | 2, 54 | 2.18 | 0.123 | |
| *PC2$_{T.myrmec}$ | HLUC | 1, 54 | 1.95 | 0.168 | |
| | Ecoregion | 2, 54 | 0.95 | 0.392 | |
| | HLUC*Ecoregion | 2, 54 | 1.16 | 0321 | |

| | | | | | |
|---|---|---|---|---|---|
| **Canopy openness** | **HLUC** | **1, 54** | **7.48** | **0.008** | **r < s** |
| | **Ecoregion** | **2, 54** | **7.81** | **0.001** | |
| | HLUC*Ecoregion | 2, 54 | 0.35 | 0.704 | |
| **Tree basal area** | HLUC | 1, 54 | 0.65 | 0.425 | |
| | **Ecoregion** | **2, 54** | **5.23** | **0.008** | |
| | HLUC*Ecoregion | 2, 54 | 1.09 | 0.342 | |
| **Shrub cover** | HLUC | 1, 54 | 1.20 | 0.277 | |
| | **Ecoregion** | **2, 54** | **9.76** | **< 0.001** | |
| | HLUC*Ecoregion | 2, 54 | 1.50 | 0.232 | |
| Other herbaceous species cover (non-myrmecochore) | HLUC | 1, 54 | 0.41 | 0.523 | |
| | Ecoregion | 2, 54 | 2.48 | 0.093 | |
| | HLUC*Ecoregion | 2, 54 | 1.17 | 0.320 | |
| Leaf habitat volume | HLUC | 1, 54 | 0.17 | 0.685 | |
| | Ecoregion | 2, 54 | 0.14 | 0.867 | |
| | HLUC*Ecoregion | 2, 54 | 0.48 | 0.621 | |
| Log habitat volume | HLUC | 1, 54 | 1.42 | 0.238 | |
| | Ecoregion | 2, 54 | 2.67 | 0.078 | |
| | HLUC*Ecoregion | 2, 54 | 2.12 | 0.130 | |
| Rock habitat volume | HLUC | 1, 54 | 0.00 | 0.950 | |
| | Ecoregion | 2, 54 | 1.43 | 0.249 | |
| | HLUC*Ecoregion | 2, 54 | 0.05 | 0.951 | |
| **Soil moisture** | HLUC | 1, 54 | 0.18 | 0.674 | |
| | **Ecoregion** | **2, 54** | **11.45** | **< 0.001** | |
| | HLUC*Ecoregion | 2, 54 | 0.32 | 0.725 | |
| **Soil temperature** | HLUC | 1, 54 | 0.14 | 0.706 | |
| | **Ecoregion** | **2, 54** | **4.42** | **0.017** | |
| | HLUC*Ecoregion | 2, 54 | 0.58 | 0.562 | |
| **Soil pH** | HLUC | 1, 54 | 0.13 | 0.717 | |
| | **Ecoregion** | **2, 54** | **4.67** | **0.013** | |
| | HLUC*Ecoregion | 2, 54 | 1.34 | 0.270 | |
| **Average air temperature** | HLUC | 1, 54 | 1.30 | 0.259 | |
| | **Ecoregion** | **2, 54** | **19.66** | **< 0.001** | |
| | HLUC*Ecoregion | 2, 54 | 1.66 | 0.200 | |

**Table S3** List of myrmecochore species with species codes and whether the species is a "significant forest indicator species" (defined as high occurrence and abundance in remnant forests compared to secondary forests) or "nonsignificant forest indicator species" (exhibited strong patterns due to HLUC but low occurrences) (see Griffiths & McGee 2018).

| Genus | Species | code | Remnant (residual) forest indicator species* or nonsignificant indicator species** (Griffiths & McGee 2018) |
|---|---|---|---|
| *Anemone* | *acutiloba* | anac | ** |
| *Anemone* | *americana* | anam | |
| *Anemone* | *quinquefolia* | anqu | |
| *Asarum* | *canadense* | asca | ** |
| *Claytonia* | *caroliniana* | clca | ** |
| *Claytonia* | *virginica* | clvi | |
| *Dicentra* | *species* | di.sp | * |
| *Dicentra* | *cucullaria* | dicu | * |
| *Erythronium* | *americanum* | eram | * |
| *Jeffersonia* | *diphylla* | jedi | |
| *Sanguinaria* | *canadensis* | saca | |
| *Thalictrum* | *thalictroides* | thth | |
| *Tiarella* | *cordifolia* | tico | * |
| *Trillium* | *erectum* | trer | * |
| *Trillium* | *grandiflorum* | trgr | * |
| *Trillium* | *species* | tr.sp | * |
| *Trillium* | *undulatum* | trun | * |
| *Uvularia* | *grandiflora* | uvgr | |
| *Uvularia* | *perfoliata* | uvpe | |
| *Uvularia* | *sessilifolia* | uvse | * |
| *Viola* | *blanda* | vibl | |
| *Viola* | *canadensis* | vica | |
| *Viola* | *pubescens* | vipu | |
| *Viola* | *rostrata* | viros | |
| *Viola* | *rotundifolia* | virot | |
| *Viola* | *sororia* | viso | |
| *Viola* | *species* | vi.sp | |
| *Viola* | *striata* | vist | |

**Table S4.** List of ant species found at the 20 forest sites from the pitfall trap surveys and seed depots. List includes species scientific name, species codes, and interaction type observed on seed depot (present, handling seeds, or carrying seeds). (Ellison et al. 2012) was used to identify ant species. Species interacting with seeds are bolded.

| Genus | Species | code | Interaction type |
|---|---|---|---|
| ***Aphaenogaster*†** | **sp.** | **aph** | **present on depot, handling, carrying** |
| *Brachymyrmex* | *depilis* | bade | |
| *Camponotus*† | *americanus* | caam | |
| *Camponotus*† | *castaneus* | caca | |
| *Camponotus*† | *chromaiodes* | cach | present on depot |
| *Camponotus*† | *nearcticus* | cane | |
| *Camponotus* | *pennsylvanicus*† | cape | present on depot |
| *Crematogaster*† | *cerasi* | chce | |
| *Formica*† | *neogagates* | foneo | present on depot |
| *Formica*† | *subaenescens* | fosubae | |
| *Formica*† | *subintegra* | fosubin | |
| *Formica* | *subsericea*† | fosubse | |
| *Hypoponera* | *punctatissima** | hypu | |
| ***Lasius*** | ***americanus*†** | **laam** | **handling** |
| *Lasius*† | *brevicornis* | labr | |
| *Lasius*† | *nearcticus* | lane | |
| *Leptothorax*† | sp1 | lepto | |
| *Myrmecina* | *americana*† | myam | |
| *Myrmica*† | *lobifrons* | mylo | |
| *Myrmica* | *punctiventris*† | mypu | present on depot |
| ***Nylanderia*** | ***flavipes**** | **nyfl** | **present on depot, handling, carrying** |
| *Nylanderia* | *parvula* | nypa | |
| *Nylanderia* | sp3 | nyla | |
| *Paratrechina* | *longicornis** | palo | |
| *Ponera* | *pennsylvanica* | pope | |
| *Prenolepis* | *imparis*† | prim | |
| *Solenopsis* | sp1 | sole | |
| *Stenamma*† | *brevicorne* | stbr | |
| *Stenamma*† | *diecki* | stdi | |
| *Stenamma*† | *impar* | stim | |
| *Stenamma* | *schmittii*† | stsc | |
| *Stigmatomma* | *pallipes* | stpa | |
| *Strumigenys* | *pulchella* | stpu | |
| *Tapinoma* | *sessile*† | tase | |
| *Temnothorax* | *curvispinosus* | tecu | present on depot |
| *Temnothorax* | *longispinosus* | telo | present on depot |
| *Temnothorax* | *schaumii* | tesc | |

* Exotic ant species

† Species or pooled genera found to interact with seeds in Ness et al. 2009.

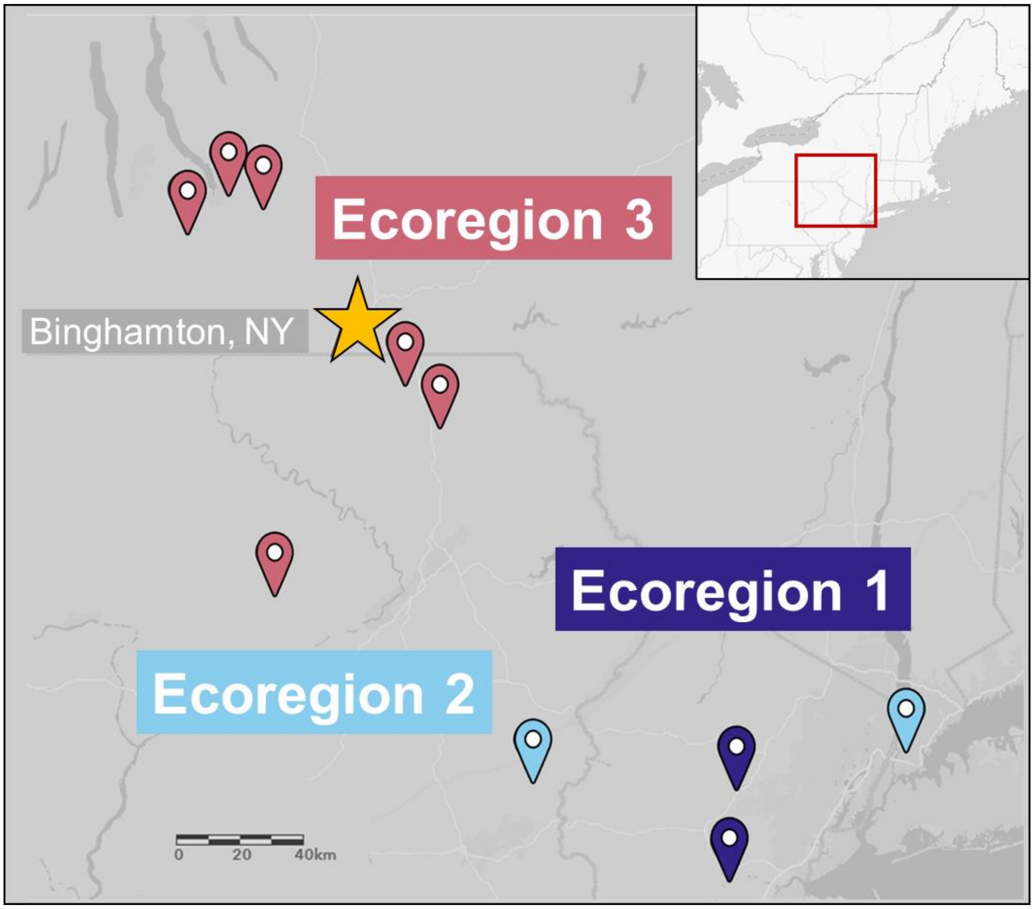


**Fig. S1** Map of study area marking pairs of remnant and secondary forested study sites in northeastern North America (n = 10 pairs, 20 sites). Different colors correspond to ecoregions, which are determined by similar ecoregions and similar dominant tree composition (see Fig. S7 and Table S1).

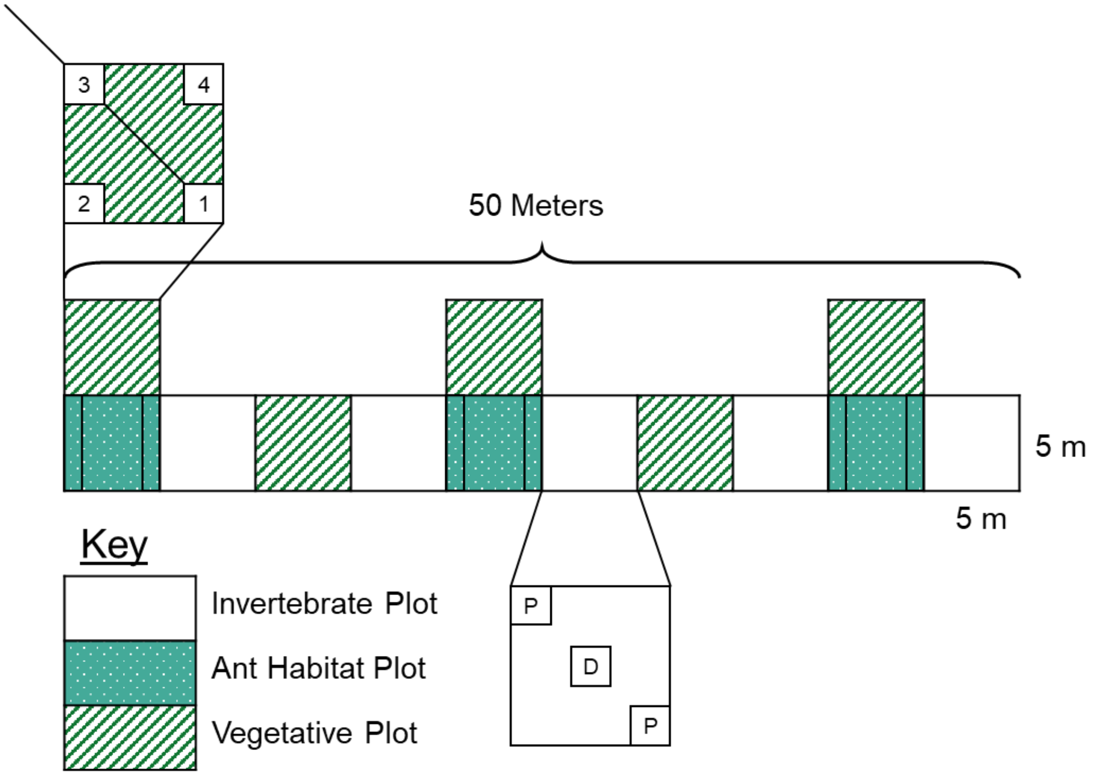


**Fig. S2** Example of one of the 3 50 m survey transect sets in the center of each forest site (n = 60). Transects consisted of 5 5x5m invertebrate plots for pitfall traps (P) and seed depots (D) (blank boxes), 3 ant habitat plots for ant habitat belt transects (solid boxes with white spots), and 5 vegetative survey plots for tree, shrub, canopy, herbaceous, and myrmecochore surveys (hatched boxes). Numbered boxes in vegetation plots indicate herbaceous survey quadrats.

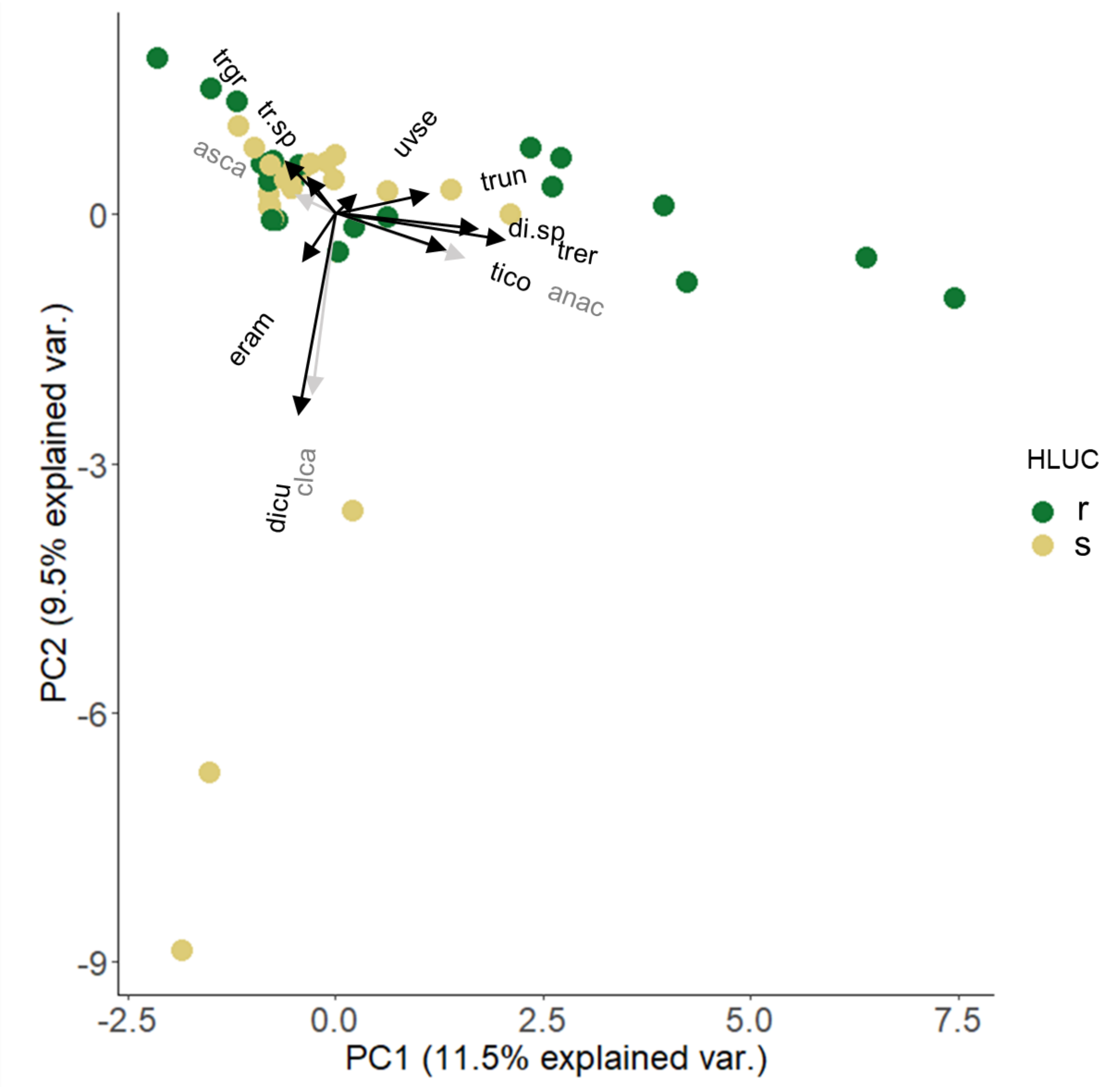


**Fig. S3** Biplot of Principle component analysis (PCA) of myrmecochore presence at the transect level. Remnant transects (r) are represented by the green symbols and secondary transects (s) the tan symbols. Remnant sites cluster more along the upper portion of $PC2_{T.myrmec}$ and positive portion of $PC1_{T.myrmec}$ and have a different myrmecochore composition compared to the majority of secondary sites which are clustered along the left side of $PC1_{T.myrmec}$. Black arrows represent indicator species and grey arrows represent nonsignificant indicator species (Griffiths & McGee 2018). Acronyms found in Table S3.

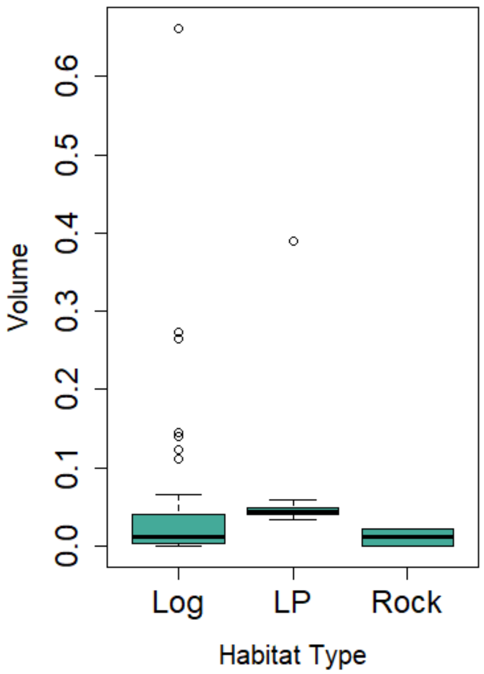


**Fig. S4** Boxplots of average volume of occupied ant habitat types (logs, leaf litter, and rocks). Thick lines in box plots represent medians, boxes represent 1st and 3rd quartiles, whiskers represent minimums and maximums, and points represent outliers. Transects and sites were combined.

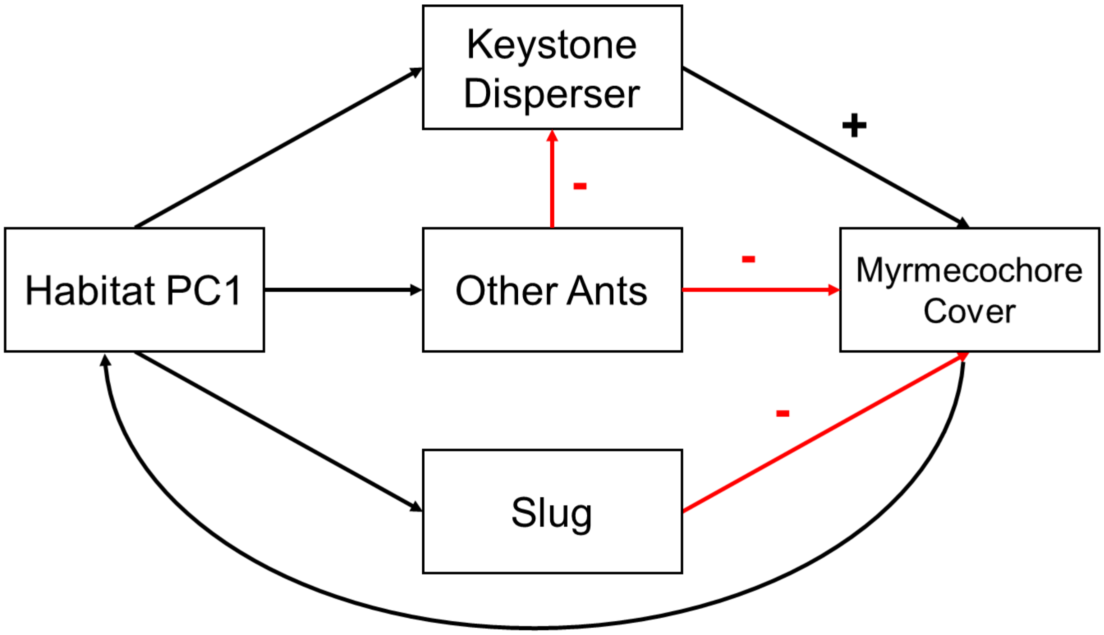


**Fig. S5** *A priori* path analysis model for seed dispersal and myrmecochore cover. Arrow directions represent assumed directionality of interactions including positive effects (black arrows) and negative effects (red arrows).

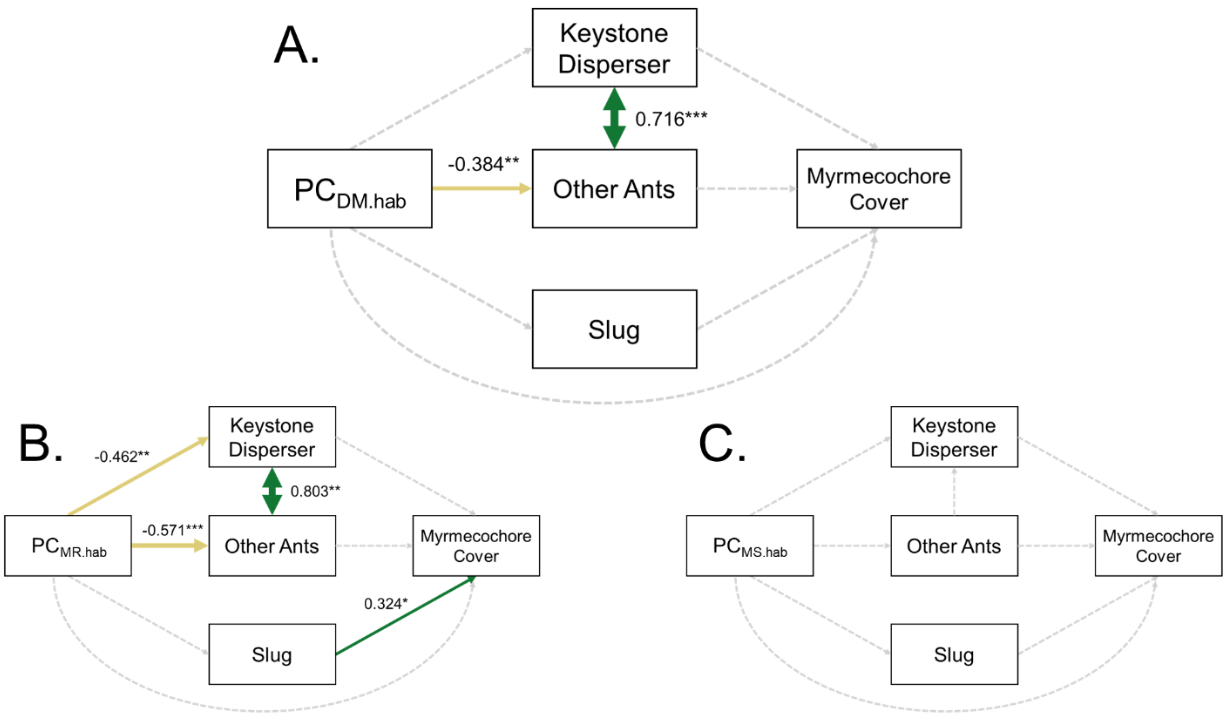


**Fig. S6** Myrmecochore cover path diagram with standardized path coefficients reported next to the arrows for A) all forests (combined) B) remnant forests and C) secondary forests. The combined path shows an effect of habitat factors ($PC_{DM.hab}$) on other ant abundance. Within remnant forests, slug abundance has a positive effect on myrmecochore cover which might be a response from increased herbaceous cover. No path coefficients in secondary forests were significant. Green and tan solid arrows indicate significant positive and negative pathways respectively. Thickness of arrows are proportional to the standardized path coefficients strength. Non-significant pathways with path coefficients less than 0.1 are given in dashed grey lines. Significance levels are indicated according to: ***$P < 0.001$; **$P < 0.01$; *$P < 0.05$.

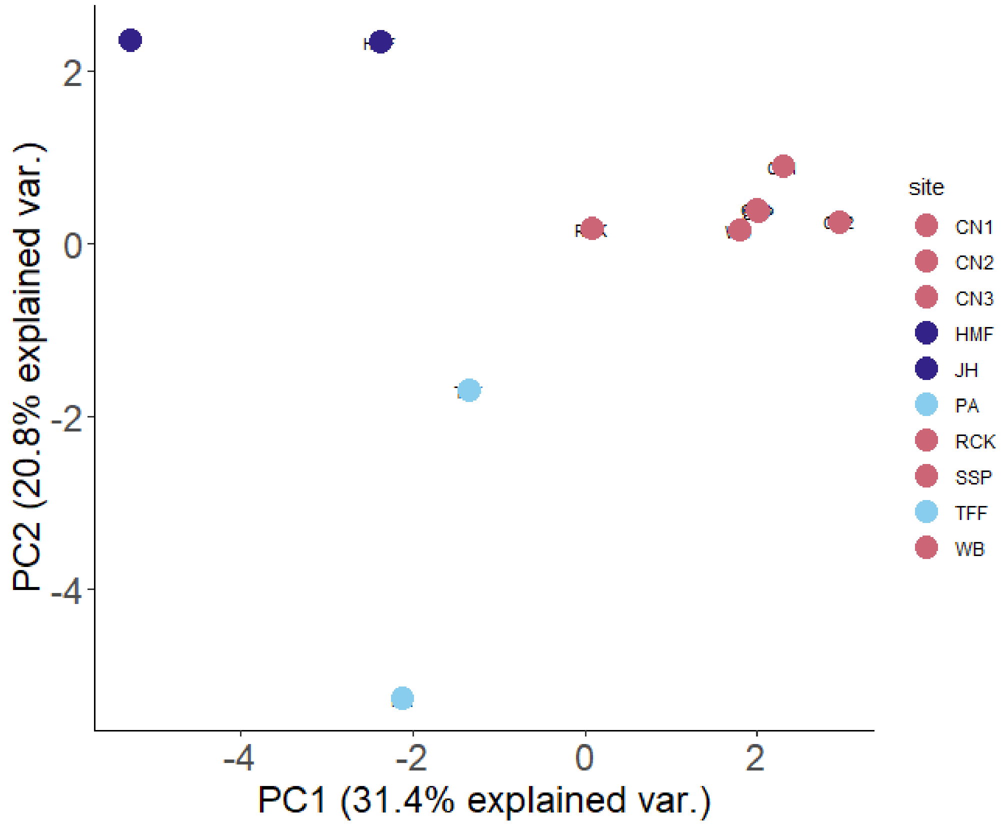


**Fig. S7** Biplot of principal component analysis (PCA) of dominant tree species at the site level.

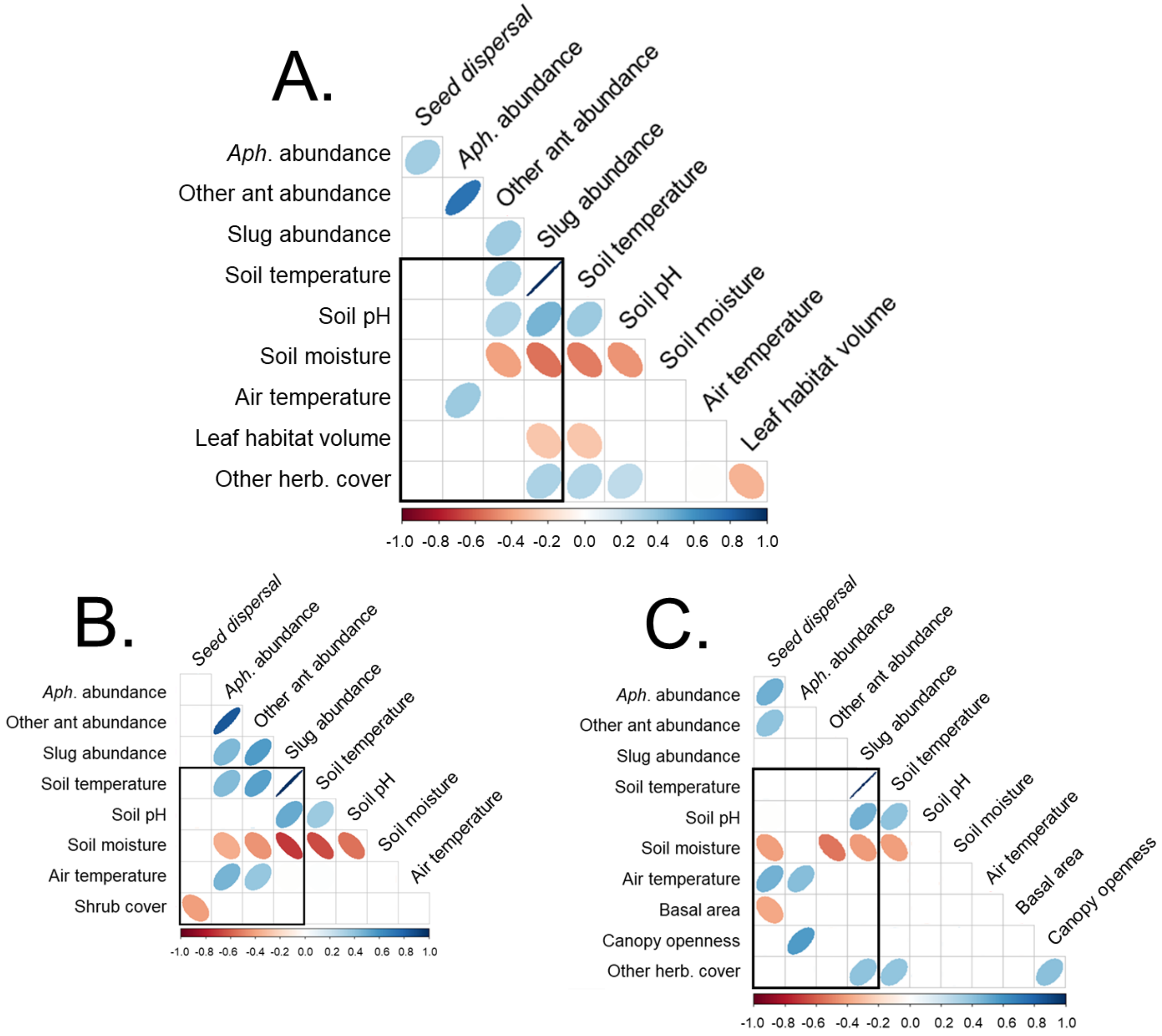


**Fig. S8** Correlation tables of habitat factors used to make composite habitat metrics for A) all forests (combined) $PC_{DM.hab}$, B) remnant forests $PC_{DR.hab}$, and C) secondary forests $PC_{DS.hab}$. Correlation analysis performed on seed dispersal response variables (seed dispersal, *Aphaenogaster* abundance, other ant abundance, and slug abundance) and habitat predictor variables (soil temperature, soil pH, soil moisture, air temperature, leaf habitat, and other herbaceous species cover, basal area, and canopy openness). Correlations among all habitat variables and seed dispersal and organisms' response variables were performed, but only significant relationships are shown. Significant correlations (indicated by ellipses) were used to make composite habitat metrics. Color and direction represent direction of correlation and width corresponds with correlation strength.

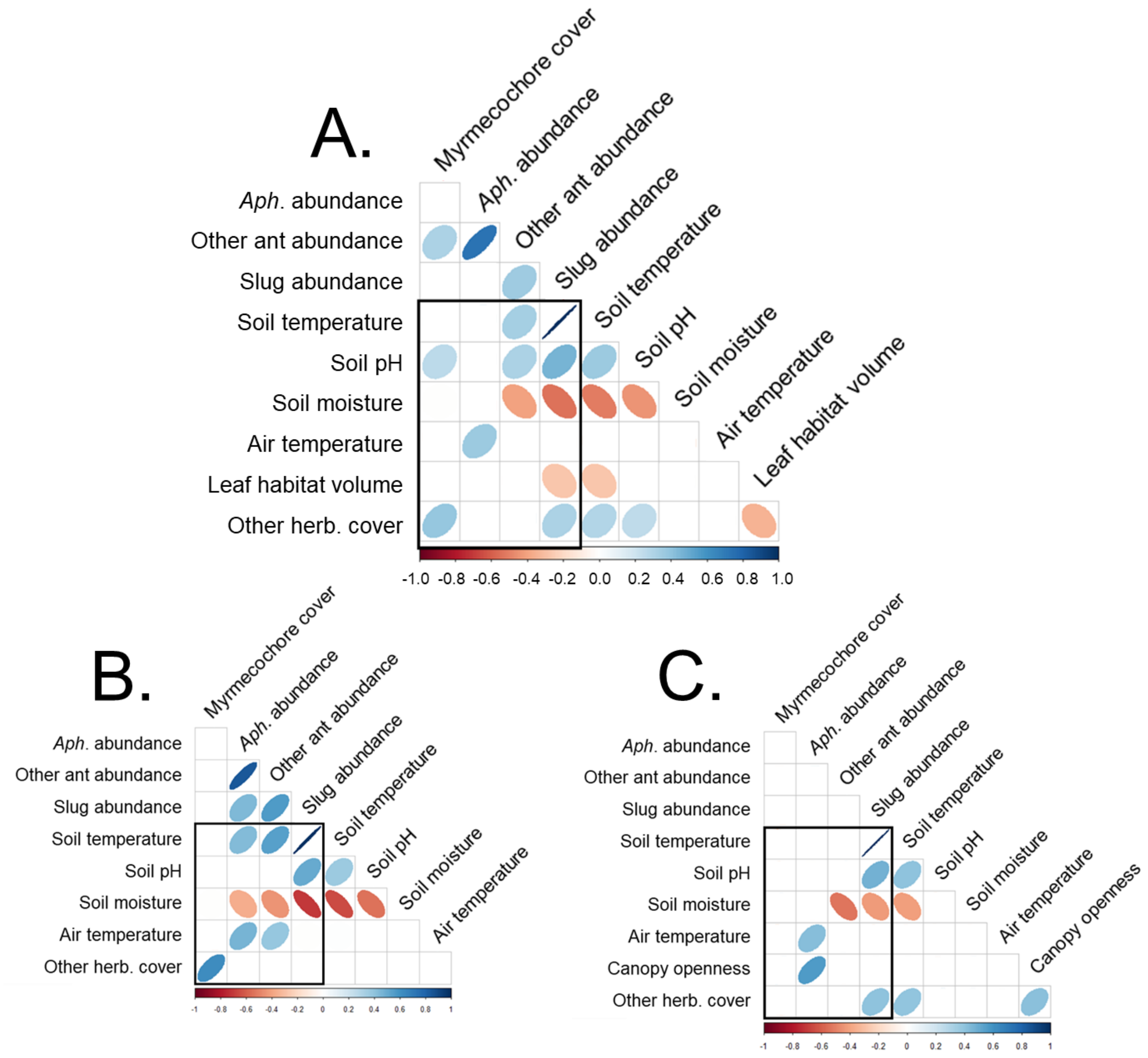


**Fig. S9** Correlation tables of habitat factors used to make composite habitat metrics for A) all forests (combined) $PC_{DM.hab}$, B) remnant forests $PC_{MR.hab}$, and C) secondary forests $PC_{MS.hab}$. Correlation analysis performed on myrmecochore response variables (myrmecochore cover, *Aphaenogaster* abundance, other ant abundance, and slug abundance) and habitat predictor variables (soil temperature, soil pH, soil moisture, air temperature, leaf habitat, and other herbaceous species cover, basal area, and canopy openness). Correlations among all habitat variables and seed dispersal and organisms' response variables were performed, but only significant relationships are shown. Significant correlations (indicated by ellipses) were used to make composite habitat metrics. Color and direction represent direction of correlation and width corresponds with correlation strength.

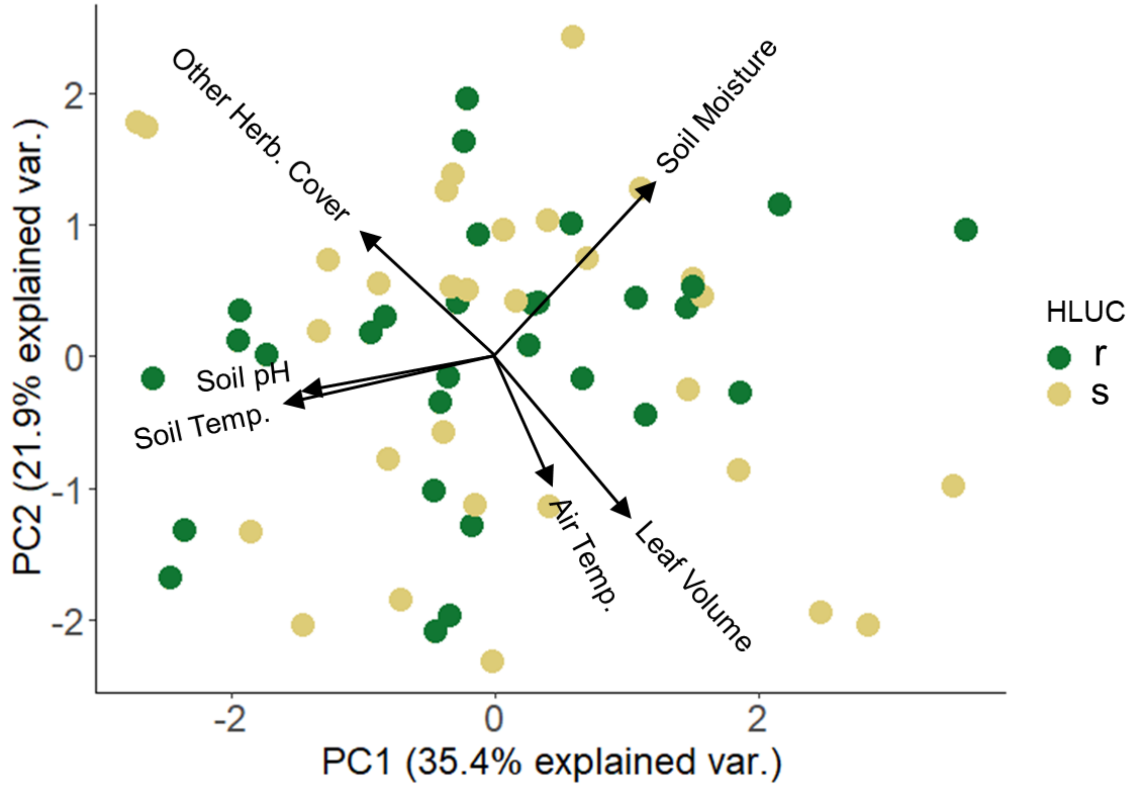


**Fig. S10** Biplot of $PC1_{DM.hab}$ and $PC2_{DM.hab}$ for seed dispersal and for myrmecochore cover combined path analyses. Points represent remnant (green) and secondary (tan) transects. $PC1_{DM.hab}$ explained 35.4% variation and $PC2_{DM.hab}$ explained 21.9% variation.
